# ROS/RNS balancing, aerobic fermentation regulation and cell cycle control – a complex early trait (‘CoV-MAC-TED’) for combating SARS-CoV-2-induced cell reprogramming

**DOI:** 10.1101/2021.06.08.447491

**Authors:** José Hélio Costa, Gunasekaran Mohanapriya, Bharadwaj Revuru, Carlos Noceda, Karine Leitão Lima Thiers, Shahid Aziz, Shivani Srivastava, Manuela Oliveira, Kapuganti Jagadis Gupta, Aprajita Kumari, Debabrata Sircar, Sarma Rajeev Kumar, Arvind Achra, Ramalingam Sathishkumar, Alok Adholeya, Birgit Arnholdt-Schmitt

## Abstract

In a perspective entitled ‘From plant survival under severe stress to anti-viral human defense’ we raised and justified the hypothesis that transcript level profiles of justified target genes established from *in vitro* somatic embryogenesis (SE) induction in plants as a reference compared to virus-induced profiles can identify differential virus signatures that link to harmful reprogramming. A standard profile of selected genes named ‘ReprogVirus’ was proposed for *in vitro*-scanning of early virus-induced reprogramming in critical primary infected cells/tissues as target trait. For data collection, the ‘ReprogVirus platform’ was initiated. This initiative aims to identify in a common effort across scientific boundaries critical virus footprints from diverse virus origins and variants as a basis for anti-viral strategy design. This approach is open for validation and extension.

In the present study, we initiated validation by experimental transcriptome data available in public domain combined with advancing plant wet lab research. We compared plant-adapted transcriptomes according to ‘RegroVirus’ complemented by alternative oxidase (AOX) genes during *de novo* programming under SE-inducing conditions with *in vitro* corona virus-induced transcriptome profiles. This approach enabled identifying a *<u>ma</u>*jor *<u>c</u>*omplex *t*rait for *<u>e</u>*arly *<u>d</u>e novo* programming during SARS-CoV-2 infection, called ‘CoV-MAC-TED’. It consists of unbalanced ROS/RNS levels, which are connected to increased aerobic fermentation that links to alpha-tubulin-based cell restructuration and progression of cell cycle.

We conclude that anti-viral/anti-SARS-CoV-2 strategies need to rigorously target ‘CoV-MAC-TED’ in primary infected nose and mouth cells through prophylactic and very early therapeutic strategies. We also discuss potential strategies in the view of the beneficial role of AOX for resilient behavior in plants.

Furthermore, following the general observation that ROS/RNS equilibration/redox homeostasis is of utmost importance at the very beginning of viral infection, we highlight that ‘de-stressing’ disease and social handling should be seen as essential part of anti-viral/anti-SARS-CoV-2 strategies.

## Introduction

Towards the tail end of 2019, a corona virus disease caused a pandemic outbreak. The virus is designated as ‘severe acute respiratory syndrome corona virus 2’ (SARS-CoV-2) and has apparently zoonotic origin. SARS-CoV-2 is a virion enveloping a single positive-stranded RNA genome, which is capable of entering into human cells by using the physiologically important, hormone-related receptor ACE2 (angiotensin converting enzyme, 2). The complex and diverse multi-systemic effects of the virus on individuals and its pandemic impact are widely discussed in comparison to two other similar viruses with endemic impacts from the *Coronaviridae* family, SARS-CoV and MERS-CoV (1–6). These similar viruses exhibit more drastic clinical impairment, but showed less human-to-human transmission rates and lower mortality (7). Also, SARS-CoV-2 has thermodynamic advantages than former Corona virus SARS-CoV through binding with higher affinity to ACE2 receptors (8).

The intense damage caused by viruses to human and the possibility of similar virus threats in the future triggered a worldwide debate and tremendous efforts have been initiated across all society levels. The global academic community aims to widen the knowledge dimensions for developing specific and wider antiviral concepts and strategies. In this context, we intend to contribute with relevant knowledge, which can be useful for developing intervention strategies well before virus spread can start to become an exponentially growing threat for the infected person and, thus, for the community.

Viruses are non-living structures. However, they are able to gain power over human cell programs and individual organism fitness due to their dominating presence. This is in conformity with knowledge on structuralism and biosemiotics in Theoretical Biology, which acknowledges the importance of relationships and signs rather than characteristics of individual components for the functioning of systems (9). In an abstract sense, virus reproduction depends on the virus-inherent ‘structural power’ in relation to other components in the system. This term is taken from peace research related to socio-economic systems, where inequality is recognized as the main driver for structural violence (10, 11). This view can also be applied for the relation of viruses to their hosts. In fact, viruses have comparatively low Gibbs energy due to their chemical compositions (12, 13). This makes their replication highly competitive and a driving force against the host cell metabolism. Virus-induced cell reprogramming favors virus replication, but demands at the same time cell reprogramming for host defense and survival. As a consequence of this conflict, any viral infection provokes struggling for commanding coordination of host cell program and this starts in the initially infected cells. Competing for bioenergy and for ‘territories’ is decisive for the success of virus reproduction and evolution.

Virus spread and its natural evolution escaped quite successfully human control. Currently, there is no awareness of any effective and simple broad anti-viral treatment available that could also be used to treat SARS-CoV-2 infections. When viral structures enter in contact with living cells, the latter cannot ignore their presence. Ignorance would be helpful and nothing harmful would happen to the host cells. However, this is way behind possible, because viral structures interact at the outer cell surface in a way that stimulates cell’s program-commanding components, such as reactive oxygen species (ROS) (14) and growth factors (15), which in turn induce changes in intracellular signaling cascades. Successful virus propagation relies on manipulated cell programming that allows abusing the host’s energy resources by help of the proper host. Since viruses are non-living structures, damage of the host is not a target but might happen as collateral circumstance (16). Therefore, virus docking to cell membranes and the subsequent membrane fusion event that enable entrance of the virus particles into living cells can be seen as ‘structural violence’. It is a sort of ‘hostile takeover’ as it is known from economy. Correspondingly, virus affects host system management and makes it act in favor of the incoming structures and risks thereby the healthy status or even life of the host cell and, finally, the overall organism (16).

Primarily virus-infected cells act as super-spreaders. In case of corona viruses, cell-cell fusion between infected and adjacent, uninfected cells were reported to form giant, multinucleated cells that allow rapid spread within infected organisms obviously even without being detected and neutralized by virus-specific antibodies (17). Massive virus replication is energy-costly, weakens and endangers the host and can cause a pandemic threat. And there is a second danger too that comes ‘secretly and as gratis’ when virus replication is not stopped during the very early phase of infection: virus coding sequences might undergo modifications due to the host’s and/or virus-driven error-prone molecular machinery. Consequently, when the rate of virus replication progresses and reaches massive propagation, the probability of virus code evolution increases and, this might enhance in turn virulence (18, 19). However, a widened diversity of virus coding structures reduces the chance that the immune system of individual organisms and of present and future networking populations are prepared to keep virus threat at low level. And thirdly, in case of SARS-CoV-2, once acquired immunity might also not last for a prolonged time (20). For these three reasons, it is of utmost importance to understand the very first steps in virus-induced cell reprogramming. This is influential to develop efficient and sustainable anti-viral concepts for confronting viral infection by administrating early or even prophylactic therapies.

In a parallel perspective paper entitled ‘From plant survival under severe stress to anti-viral human defense’ we proposed using a standard profile of selected genes (‘ReprogVirus’) to trace early footprints of wide varieties of viruses under standardized *in-vitro* conditions (21). We raised and justified the hypothesis that transcript level profiles of justified target genes established from *in vitro* somatic embryogenesis (SE) induction in plants as a reference compared to virus-induced profiles can identify differential virus signatures that link to harmful reprogramming. This interdisciplinary approach was explained in details, including especially the use of SE induction as an efficient experimental tool to identify markers for early reprogramming and resilience and the focus on alternative oxidase (AOX). Here, we follow strictly that approach in order to initiate scientific validation of the underlying hypothesis and enable progressing in a common effort towards the wider perspective for anti-viral strategy-design.

## Background

### Virus captures host cell signaling and metabolism

Viral infections causing respiratory complications are known to change host cell redox homeostasis, which involves balancing ROS/RNS as a critical event (22). RNA viruses were suggested to utilize oxidative stress during infection to control genome RNA capping and genome replication (14, 23, 24). In airway epithelial cells, virus-induced ROS was found to originate from diverse oxidase activities, including NADPH oxidases, dual oxidase, and xanthine oxidase (reviewed in 22). Similar to plant superoxide dismutase (SOD), animal SOD plays a central role as an oxidative stress indicator and also as an anti-oxidative stress defender along with a set of other ROS scavenging enzymes, such as, catalase, GPX (glutathione peroxidase) and GSR (glutathione reductase) was found to be induced upon viral infection (25) and also highlighted in stem cell research as an indicator for cell reprogramming (26). Likewise, nitric oxide (NO) was found to be involved in virus replication (27, 28). In humans, NO production during viral infection depends on nitric oxide synthase (*NOS*). Inducible NOS (NOS2) produced much higher amounts of NO for a prolonged duration as compared to constitutively expressed neuronal (*NOS1*), endothelial (*eNOS* or *NOS3*) and mitochondrial *NOS* (29). Biogenesis of higher levels of NO can suppress type 1 helper T-cell-dependent immune responses, which can impair type 2 helper T-cell-biased immunological host responses (29). eNOS is mostly present in endothelial cells and its functionality can be restored with renin- and angiotensin-converting enzyme-inhibitors or angiotensin receptor blockers, both commonly used to regulate blood pressure in hypertensive patients (30).

Several metabolic changes have been reported in virus-infected animal and/or plant host cells that were related to central carbon metabolism: e.g. (a) increased rate of glycolysis linked to pools of nucleotides and amino acids essential for replication (31, 32), (b) differential mTOR (mammalian target of rapamycin) pathway regulation and intracellular calcium signaling linked to enhanced Krebs cycle, (c) down-regulation of glycolysis that severely affected viral infection (33–35), (d) encoding mitochondria-related proteins that disturbed normal functioning of mitochondria (36), (e) increased production of lactic acid from glucose pumped out of cell (37), (f) knock-down of *ADH* (*Alcohol dehydrogenase*) and pyruvate decarboxylase that diminished virus replication rates (38), and, (g) interplay between glycolysis and fermentation that is suggested to serve metabolome channeling for virus replication highlighted as paradigm change (39). Bojkova et al. (40) observed that blocking of glycolysis resulted in prevention of SARS-CoV-2 replication in infected cells. Lactate present in the circulating system had been identified besides interleukin-6 as independent prognostic factors for disease progression in a relatively small clinical data collection. In addition, increased lactate was found to impair antiviral immunity (41). Subsequent to the appearance of symptoms, disease severity was classified by monitoring disease progression during 24h to 48h and beyond this duration of disease progression, lactate was not found to be useful to predict fatality (survivor vs non-survivor) (42). In primary lung epithelial cells validated by biopsies of COVID-19 patients, SARS-CoV-2 induced oxidative stress due to mitochondrial dysfunction (32). In these studies, endoplasmic stress symptoms were related to modulation of lipid metabolism.

Transcriptional responses in primary lung epithelial cells and data from biopsies were shown to be predominantly of metabolic nature (59% to 65% of all differentially expressed genes). They indicated upregulation of glycolysis and dysregulation of citric acid cycle, which was mediated by the transcription factors *NF-kB* and *RELA*. In patients, elevated blood glucose was found to be a risk factor independently from diabetes. SARS-CoV-2 infection was associated to marked increase in intracellular glucose and transcriptional modulation in glucose metabolism along with elevated lactate levels (32). Several authors report that in virus research transcriptome data and a focus on gene sets revealed to be critically relevant for cell and organism performance (43–47).

### Host microtubule assembly is critical for virus entry, replication and spread

Microtubules (MT) interfere with viral infection process right from the very start. Thus, the aspects related to regulation of biosynthesis and organization of the MT assembly determines critically the frame for reprogramming and the acquiring of basal defense of the host cells. Virus attach and enter at specific sites of human host cells in relation to the spatial organization of surface receptors and entry factors, which is partially controlled by MT and their polarization (58). MT-driven transport within the cells by dynein and kinesin family of motors has been shown to have crucial role in virus replication and spread (59). Interaction of alpha-corona virus spike (S) proteins with tubulin has been shown to support S protein transport and incorporation into virus particles (60). Virus can promote MT polymerization, stabilization or disruption and actin-MT crosstalk depending on the cellular status with regard to the stage in viral infection. Receptor engagement by viruses’ influences MT dynamics facilitating viral infection during the early stages of infection. MTs or MT-like proteinaceous filaments are found to be highly relevant for infections by diverse viral classes and across kingdoms and species (58). α-Tubulin has been shown to be involved in regulating cell death and program decisions linked to cell cycle activation (61, 62). Enzymes catalyzing post-translational MT modifications are realized to be suitable targets for drug development for combating viral infection (62).

### Viruses can influence host cell cycle in favor of their own replication

Viruses can affect host cell cycle regulation in favor of viral replication, which might seriously affect host cell physiology with impacts on pathogenesis (48–50). Viruses were shown to influence different stages of the host cell cycle resulting in either progression or arrest of cell cycle with the involvement of homologous proteins (49, 51). Cellular levels of phosphorylated retinoblastoma tumor suppressor protein (pRb) were found to be critical for the progression of cell division to S phase (52). Hyperphosphorylation of Rb allows activation of E2F family of transcription factors, resulting in the transcription of genes associated with S phase of the cell cycle (53). In Rb-deficient mouse embryos, mutation of E2F1 was found to markedly suppress apoptosis and also entry of cells into S-phase resulting in prolonged cell survival (54). Impressively, TOR-suppression in plants either by silencing or by inhibitor treatment (AZD8055) was shown to reduce virus replication, leading to conferring virus resistance in the host plant or even to total elimination of viral infection (reviewed in 55). Corona viruses have been shown to notably arrest cells in G0/G1 stage of cell cycle. Enrichment of coronavirus-infected cells had also been found in G2/M stage (49). SARS-CoV is known to produce 3b and 7a non-structural proteins, which together decrease level of cyclin D3 and dephosphorylation of pRb in order to arrest cell cycle at the S phase (56). SARS-CoV-induced synthesis of protein 3a was reported to arrest cell cycle at the G1 stage via the operation of the earlier mentioned pathways (57). Recently, Laise et al. (50) identified 12h, 24h, and 48h signatures from calu-3 lung adenocarcinoma cells infected with SARS-CoV, which included genes related to cell cycle progression, viz., E2F and mTOR.

### Host microbiome can influence viral infection progress

Host microbiomes are individual specific. They can affect viral infection either negatively and positively (63, 64). Host microbiome can increase virion stability, provide DNA replication machinery, stimulate the lytic phase of virus and counteract host immune responses. Further, microbiota can even enhance viral genetic recombination through adhering host cells and facilitating thereby infection of two or more viruses in a single cell (65). In contrast, probiotic bacteria, such as *Bifidobacterium* and *Lactobacillus,* increased cellular biosynthesis of cytokines and interleukins upon viral infection (66, 67). Additionally, *Lactobacillus vaginalis* inhibited infection through human immunodeficiency virus (HIV) by producing lactic acid and maintaining acidic environment in vagina (68). Vaginal lactic acid can profoundly increase biosynthesis of anti-inflammatory cytokines, which was shown to help preventing infection by Herpes virus (69). These observations demonstrate that it can be important to better understand the interaction of microbiota with virus-induced early reprogramming in target cells and to apply this knowledge when prophylactic and early therapies are required.

## Material and Methods

### Gene expression analyses of RNA-seq data from *Arabidopsis thaliana* and virus-infected human cells

Genes for expression analyses correspond to standard profile of genes selected for studying virus-induced early reprogramming (named ‘ReprogVirus’) (21, in press). For plant material, ‘ReprogVirus’ was complemented by alternative oxidase (*AOX*) and genes of plant secondary metabolism. In summary, the selected genes related to ROS/RNS equilibration (*AOX*, *ADH2*/*ADH5*), anti-oxidant activities (*SOD, Cat*, *GPX*, *GSR*, plant-*PAL*, *CHS*, *C3H*, *CAD*), NO production (plant-*NR*, *NOS*), glycolysis (*Hexokinase*, *PFK-M*, *GAPDH*, *enolase*, *pyruvate kinase*), *G6PDH*, *MDH1/2*, fermentation (*LDH*, *ADH1*), structural cell organization (*alpha-tubulin*, *actin*), energy status-signaling (*SNRK*), cell cycle regulation (*TOR/mTOR*, *E2F1-3*, *E2F5*) and regulation of apoptosis/cell death (*BAG*, *Meta-CASP*, *Caspase In*, *Caspase Ex*, *Bcl-xL*). In the case of virus-infection trials, a set of additional genes from the immune system (*IRF9* and *IRF3*) and the two transcription factors NF-KB1 and NF-KB-RELA were included. For SARS-CoV-2related profiles, we added receptor ACE2 and the priming protease *TMPRSS2*. Additionally, we searched transcriptomes of virus-infected cells for the melatonin synthesis-related gene *ASMT* (N-acetylserotonin O-methyltransferase). *ASMT* is involved in melatonin synthesis in human cells (70).

Gene expression was evaluated in specific RNA-seq experiments, which were collected from Sequence Read Archive (SRA) database of GenBank, NCBI. Expression analyses were performed from *Arabidopsis* and virus infected human cells. Experimental and projects details were given in Supplementary Table S11. Specific regions (3’ end) of each cDNA were aligned against RNA-seq experiments using mega BLASTn tool (71) to obtain the mapped reads according to Saraiva et al. (72). Specific parameters of mega BLASTn as word size were adjusted to allow specific read detection of each gene. The mapped reads were also verified using the Magic-Blast (73), a more recent tool with accurate features for RNA-seq data. The number of mapped reads (on each gene/experiment) was normalized using the RPKM (Reads Per Kilo base of transcript per Million of mapped reads) method (74). The following equation was applied: RPKM = (number of mapped reads X 10^9^) / (number of sequences in each database X number of nucleotides of each gene). According to the mined datasets, transcript levels of *Arabidopsis* seedlings consisted of the values of three biological replicates. All viral infection experiments had three biological replicates, except RSV infection to A459 cells, which had only two biological replicates. The number of reflected technical replicates varied in the RSV datasets between one and three.

### All selected genes are listed below

For *Arabidopsis*: total *AOX* (Alternative oxidase) [*AOX1a* (AT3G22370.1); *AOX1b* (AT3G22360.1); *AOX1c* (AT3G27620.1); *AOX1d* (AT1G32350.1); *AOX2* (AT5G64210.1)], *LDH* (Lactate dehydrogenase) (AT4G17260.1), *ADH1* (Alcohol dehydrogenase) (AT1G77120.1), Total enolase [*cyt-ENO1* (AT2G29560.1); *cyt-ENO2* (Cytosolic-enolase) (AT2G36530.1); *plast-ENO1* (Plastidial-enolase) (AT1G74030.1)], *cyt-Fe-SOD1* (Cytosolic-iron-Superoxide dismutase 1) (AT4G25100.1), *mt_Mn-SOD1* (mitochondrial manganese superoxide dismutase 1) (AT3G10920.1), *CAT3* (catalase), (AT1G20620.1), Total *GPX* (Gluthatione peroxidase) [*GPX1* (AT2G25080.1); *GPX2* (AT2G31570.1); *GPX3* (AT2G43350.1); *GPX4* (AT2G48150.1); *GPX5* (AT3G63080.1); *GPX6* (AT4G11600.1); *GPX7* (AT4G31870.1); *GPX8* (AT1G63460.1)], *Cyt-GSR1* (Gluthatione reductase) (AT3G24170.1), total *PAL* (Phenylalanine ammonia lyase) [*PAL1* (AT2G37040.1); *PAL2* (AT3G53260.1); *PAL3* (AT5G04230.1); *PAL4* (AT3G10340.1)], total *CHS* (Chalcone synthase) [*CHS-A* (AT1G02050.1); *CHS-B* (AT4G34850.1); *CHS-C* (AT4G00040.1); *CHS-D* (AT5G13930.1)], *C3H* (p-coumarate 3-hdroxylase) (AT2G40890.1), *CAD* (cinnamyl alcohol dehydrogenase) [*CAD4* (At3g19450); *CAD5* (At4g34230); *CAD7* (At4g37980); *CAD8* (At4g37990)], total *NR* (nitrate reductase) [*NIA1* (AT1G77760.1); *NIA2* (AT1G37130.1)], *ADH2* (AT5G43940.2), total *ACT* (actin) [*ACT1* (AT2G37620.1); *ACT2* (AT3G18780.1); *ACT3* (AT3G53750.1); *ACT4* (AT5G59370.1); *ACT7* (AT5G09810.1); *ACT8* (AT1G49240.1); *ACT9* (AT2G42090.1); *ACT11* (AT3G12110.1); *ACT12* (AT3G46520.1); *ACT* (AT2G42170.1)], total alpha-tubulin [*TUA1* (AT1G64740.1); *TUA2* (AT1G50010.1); *TUA3* (AT5G19770.1); *TUA4* (AT1G04820.1); *TUA5* (AT5G19780.1); *TUA6* (AT4G14960.2)], *SNRK* (sucrose non-fermenting related kinase) [*KIN10* (AT3G01090.1); *KIN11* (AT3G29160.1)], *TOR* (target of rapamycin) (AT1G50030.1), *E2F* [E2F1 (AT5G22220.2); *E2F3* (AT2G36010.1); *E2F5* (AT1G47870.1)], total *BAG* (Bcl-2 associated gene) [*BAG1* (AT5G52060.1); *BAG2* (AT5G62100.1); *BAG3* (AT5G07220.1); *BAG4* (AT3G51780.1); *BAG5* (AT1G12060.1); *BAG6* (AT2G46240.1); *BAG7* (AT562390.1)], Meta-caspase (*MC-1* (AT1G02170.1); *MC-2* (AT4G25110.1) *MC-3* (AT5G64240.1); *MC-4* (AT1G79340.1); *MC-5* (AT1G79330.1); *MC-6* (AT1G79320.1); *MC-7* (AT1G79310.1); *MC-8* (AT1G16420.1); MC-9 (AT5G04200.1)], *ASMT* (AT4G35160).

For *Homo sapiens*: total *LDH* [*LDH-A* (NM_005566.4); *LDH-B* (NM_002300.8); *LDH-C* (NM_002301.4); *LDH-AL6A* (NM_144972.5); *LDH-AL6B* (NM_033195.3)], *SOD1* (NM_000454.5), *SOD2* (M36693.1), *Catalase* (NM_001752.4), total *GPX* [*GPX-1* (NM_000581.4); *GPX-2* (NM_002083.4); *GPX-3* (NM_002084.5); *GPX-4* (NM_002085.5); *GPX-5* (NM_001509.3); *GPX-6* (NM_182701.1); *GPX-7* (NM_015696.5); *GPX-8* (NM_001008397.4)], *GSR* (NM_000637.5), *NOS1* (nitric oxide synthase) (NM_000620.5), *NOS2* (NM_000625.4), *NOS3* (NM_000603.5), *ADH5* (NM_000671.4), *Hexokinase* [*HK1* (NM_000188.3); *HK2* (NM_000189.5); *HK3* (NM_002115.3)], *PFK-M* (NM_001166686.2), *GAPDH* (NM_002046.7), *Enolase* [*Eno1* (NM_001428.5); *Eno2* (NM_001975.3); *Eno3* (NM_001976.5)], *Pyruvate kinase* [*PKLR* (XM_006711386.4); *PKM* (NM_002654.6)], *G6PDH* (NM_000402.4), *MDH1* (NM_005917.4), *MDH2* (NM_005918.4), *SNRK* (NM_017719.5), *mTOR* (NM_004958.4), *E2F1* (NM_005225.3), *Actin* [*ACT-A1* (NM_001100.4); *ACT-B* (NM_001101.5); *ACT-G1* (NM_001199954.2)], *IRF9* (NM_006084.5), *IRF3* (NM_001571.6), *NF-KB1* (NM_003998.4), *NF-KB-RELA* (NM_021975.4), *Caspase in* [*CASP8* (NM_001228.4); *CASP9* (NM_001229.5); *CASP10* (NM_032977.4)], *Caspase ex* [*CASP3* (NM_004346.4); *CASP6* (NM_001226.4); *CASP7* (NM_001227.5)], *BCL-xL* (Z23115.1), *ACE2* (NM_001371415.1), *TMPRSS2* (NM_001135099.1), *ASMT* (NM_001171038.2).

### Induction of somatic embryogenesis in *Daucus carota* L

Carrot seeds (cv. Kuroda) were surface-sterilized with 75% ethanol for 1 min and 4% sodium hypochlorite solution for 20 min. Later, seeds were washed thrice with sterile water and dried on filter paper under laminar air flow. Dried seeds were collected into sterile screw-cap tubes and stored in dark at room temperature. Surface sterilized seeds were inoculated on B5 solid medium supplemented with 0.5 mg l^-1^ 2,4-D according to Mohanapriya et al. (75), also supplemented with 0% and 2% sucrose. Plates were incubated in culture room at 22-25°C with 16h/8h photoperiod. Triplicates of 40 seeds per plate were maintained for each treatment. At 0, 6, 12, 24, 48 and 72 hours after inoculation (HAI) numbers of seeds with induced callus were recorded and samples were collected simultaneously for ADH assay. The embryogenic nature of calli was routinely evaluated by microscope till 45 days after inoculation. With 0% sucrose, non-embryogenic calli were induced, while in presence of 2% sucrose, embryogenic calli were obtained. In this system, the efficiency of SE induction can be optimized by increasing sucrose to 3%, that induces a significantly higher number of calli leading to SE when compared to 2% sucrose (unpublished data).

For studying the influence of the *AOX*, inhibitor salicyl hydroxamic acid (SHAM) and its interaction with sucrose on the morphogenetic effect of auxin-treatment, seeds were inoculated on 0.5 mg l^-1^ 2,4-D supplemented solid B5 medium at 0% and 3% sucrose along with 10 mM SHAM (dissolved in 50% DMSO and sterilized by using 0.22 µm filters). Seeds were harvested at 0, 6 and 12HAI, and 10 days after inoculation, and then shifted into SHAM-free media. Emergence of calli were recorded at every 24h intervals until 240 HAI (24, 48, 72, 96, 120, 144, 168, 192, 216, and 240 HAI). Experiments of callus differentiation consisted in two sets of 40 seeds each per treatment, and ADH measures were performed with duplicated pools of 40 seeds, according to the description of the experiment.

### Alcohol dehydrogenase (ADH) assay performed during SE induction from carrot seeds

Forty seeds were collected at 0, 12, 24, 30, 48, and 72 HAI grown on 0.5 mg l^-1^ 2,4-D supplemented solid B5 medium. They were bulked and ground into powder in a sterile mortar and pestle by using liquid nitrogen. Fine powder was extracted using 1 mL of 1X phosphate saline buffer (pH 7.2) and centrifuged for 30 min at 10,000 rpm at 4^°^C. The obtained supernatant was used as crude protein extract for enzyme assay. The assay was performed according to the protocol given in Kagi and Vallee (76). Briefly, the reaction mixture consisted of 1.3 ml 50 mM sodium phosphate buffer (pH 8.8), 0.1 ml 95% ethanol, 1.5 ml 5 mM *ß*-nicotinamide adenine dinucleotide (*ß*-NAD^+^) and 0.1 ml of crude protein extract. This mixture was immediately inversed several times for homogenization and absorbance at 340 nm was recorded from 0 to 6 min. Alcohol dehydrogenase from *Saccharomyces cerevisiae* was used as positive control. Active enzyme units (IU - international units) were calculated according to the following formula:

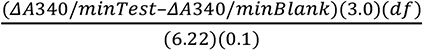

3.0 = Total volume (ml) of assay

df = Dilution factor

6.22 = millimolar extinction coefficient of reduced ß-nicotinamide adenine dinucleotide (ß-NAD^+^) at 340 nm

0.1 = Volume (ml) of crude extract used

In **Figure 1A.2**, the calculated IU of ADH were multiplied by a factor (5) for better visualization of the relationship between ADH (IU) and number of seeds with showing callus.

**Figure 1:**
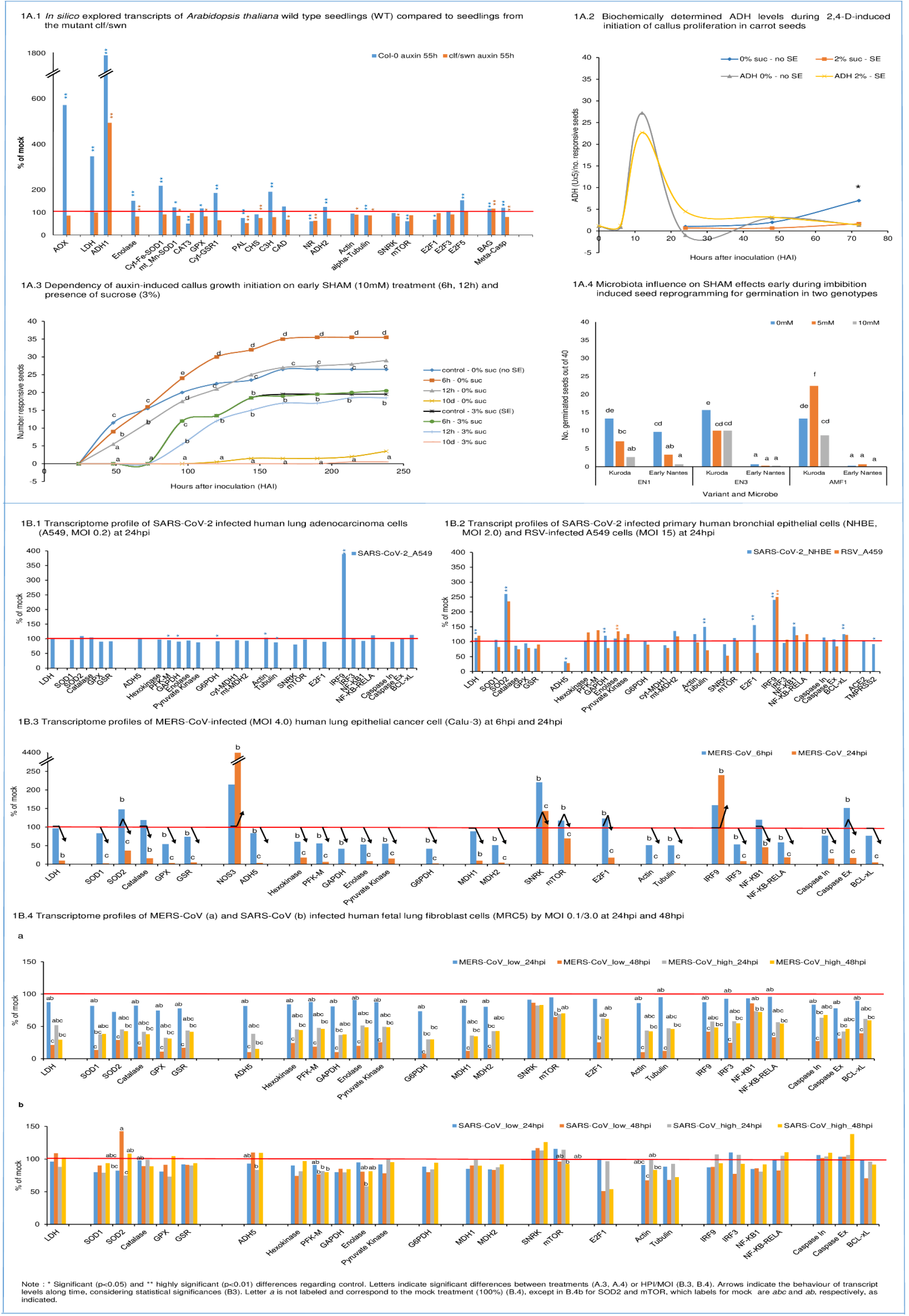
Functional transcriptome profiles at early reprogramming. Figure 1A.1: *In silico* explored transcripts of *Arabidopsis thaliana* wild type (WT) seedlings compared to seedlings from the mutant clf/swn. **Results:** (**1**) At 55h of auxin treatment, the mutant did not show signs of oxidative stress: none of the selected transcripts involved in oxidative stress regulation were increased against the control and *AOX* remained close to control level. In contrast, the according transcript profile of WT seedlings indicated at that time high oxidative stress. *AOX* shows strongly enhanced transcript accumulation against the control (5.71 folds, significant) along increased complex transcript levels of several anti-oxidants (*SOD*, *GPX*, *GSR1*, *CH3*, *CAD*). (**2**) WT displays at 55h of auxin treatment strongly decreased *NOS1*- and also significantly reduced *NR* - transcript levels, which together indicate down-regulation of NO production. Significantly decreased *NOS1* transcript accumulation was also observed for the mutant though less pronounced. *NR* transcript level was indicated, but non-significant. No significant differences to the controls were observed for *ADH2*/*GSNOR* in both variants. (**3**) In both variants, we observed after 55h of auxin-treatment similarly reduced transcript levels for alpha-tubulin (significant). For actin, the mutant shows the same significant reduction in transcript levels as for tubulin, while in WT a reduction for actin is only indicated (non-significant). In WT, increased transcript levels for *E2F5* (cell cycle suppressor gene) associated to reduced levels of cell cycle activator *E2F1* (both significant) and equal-to-control levels for *E2F3* (activator) at almost unchanged SNRK but lowered TOR transcript levels (significant). All together, these results indicate cell cycle suppression for WT at this early stage of reprogramming. In contrast, the mutant showed only slightly decreased *TOR* and *E2F1/3* transcript levels (all non-significant) and no increase in *E2F5* transcripts, but significantly reduced *SNRK* transcripts. Collectively, these results signal that induction of cell cycle progression towards embryogenic callus growth were for the mutant already more advanced than non-embryogenic callus growth induction for WT. However, in both cases, cell cycle arrest was indicated for that time point. (**4**) In WT, fermentation-related *ADH1* and *LDH* gene transcripts were strongly enhanced (17.8 folds and 3.5 folds, both significant), which associated to increased glycolysis presented by *Enolase* transcripts (1.5 fold significant). In the mutant, the only profile components that increased against the respective mock control were *ADH1* and *BAG* (both significant). However, at 55h of auxin-treatment this increase in *ADH1* in the mutant (4.9 folds of mock) was clearly and significantly less than the increase observed for WT (17.8 folds). In **Figure 1** and Supplementary table S1-S5 (both in Supplementary files), it can be seen that the mutant showed in the control higher transcript levels than WT control for *ADH1* (3.9 folds of WT, significant), *LDH* (2.4 folds of WT, non-significant) and *Enolase* (1.3 folds of WT, non-significant). However, after auxin treatment, the increased absolute *ADH1* transcript levels had been similar between both variants (1.1 fold of WT), whereas *LDH* levels remained basal and linked to decreased enolase. Higher levels of transcripts related to aerobic fermentation in the mutant controls were connected to higher *AOX* transcript levels (1.37 folds, significant). However, during auxin treatment *AOX1* transcript levels were strongly increased in WT (significant), indicating a stress situation, while no further increased *AOX1* transcript accumulation was observed in the mutant. This demonstrates that increased *AOX* transcript levels in the WT during auxin-induced reprogramming corresponded to acute metabolic requirements that were not given in the mutant. Overall, these results point to a higher basal capacity of the mutant to limit aerobic fermentation as relevant factor for the mutant’s generally higher efficiency for auxin-inducible reprogramming and an associated role for AOX. This capacity was associated to already higher transcript levels of *AOX* in the mutant control. (**5**) A more advanced stage of reprogramming for the mutant is also obvious by looking to apoptosis- or, in general, cell death-related genes, such as *BAG* (Bcl-2-related genes) and *Meta-caspase*. *BAG* transcript levels were similarly enhanced in both variants (significant). However, while meta-caspase gene transcripts in WT were increased to about the same level as for *BAG* (significant), in the mutant, *Meta-caspase* transcript levels were down-regulated (significant). This observation indicates that (a) activation of the complex cell death-related regulatory system formed part of the reprogramming process and that (b) cell death-promoting enzymes were down-regulated during the later phase of reprogramming. **Figure 1A.2: Biochemically determined ADH levels during 2,4-D-induced initiation of callus cell proliferation in carrot seeds** **Results:** Sucrose postponed initiation of callus growth from second day onwards (significant at 72HAI). The initial arrest of callus growth associated to a lower ADH peak at 12h. Although the discrepancy in ADH peaks here shown was non-significant, we had observed in further trials that higher sucrose supply (3%) further reduced this ADH peak level at 12h (84, preprint). In conclusion, these results together with the findings in Fig. 1A.1 confirm that reprogramming is, in general, linked to temporarily up-regulated ADH. It indicates a general role for early regulated aerobic fermentation in reprogramming. In the three experimental systems described in the two figures, which include also different plant species and genotypes, cell proliferation was suppressed at the earliest stage during *de novo* reprogramming and this was independently on later cell destinies. These results also show that sucrose can be a critical factor for fermentation-related reprogramming during its early phase. **Figure 1A.3: Dependency of auxin-induced callus growth initiation on early SHAM (10mM) treatment (6h, 12h) and presence of sucrose (3%)** **Results:** SHAM significantly affected callus growth initiation at 0% sucrose. At 48HAI and 72HAI, it could be observed that SHAM suppressed emergence of callus growth with time of treatment duration (6HAI, 12HAI to its permanent presence). In contrast, from 96HAI onwards, a short initial pulse of SHAM (6h) was sufficient to increase rate of callus emergence (significant). On the other hand, if SHAM supply was prolonged to 12hrs, callus emergence rate was similar to the control. However, in the presence of external sucrose (3%), short SHAM pulses of 6 or 12h did not affect callus initiation. Callus emergence was postponed to about the same degree as observed at 3% sucrose without SHAM and the growth curves were similar. This indicates that oxidative stress regulation and AOX involvement interact with sucrose. When SHAM was present during all 10 days of the trial, callus growth was suppressed in control and sucrose-containing media, and thus, also embryogenic development was suppressed. Overall, these results point to a superimposed role of fine-tuned oxidative metabolism/redox status regulation for hormone-dependent metabolic reprogramming during early induction and also during later growth initiation and highlight its interaction with sucrose. These results also validate central AOX functionality for efficient cell reprogramming under stress, which is highly relevant for breeding on plant resilience (75, 87). **Figure 1A.4: Microbiota influence on SHAM effects early during imbibition-induced seed reprogramming for germination in two genotypes** **Results:** SHAM had differential effects on root emergence monitored at 48 hours after imbibition dependent on microbiota treatment. In cultivar Kuroda, 5mM SHAM together with AMF improved germination, whereas treatment of 5mM SHAM together with EN1 and EN3 reduced root emergence. While SHAM effects had been dose-dependent for EN1 and AMF, under EN3 + AMF treatment, the higher concentration of SHAM did not lead to less germination. The cultivar Early Nantes is germinating later and only under EN1 treatment, SHAM reduced germination in this genotype. We identified main effects for all three factors (plant genotype, microorganism and SHAM concentration), and interactions for all factor combinations. These results point to the general importance of the holobiont nature of cells and individual organisms when considering oxidative metabolism/redox status regulation. They also support that genotype-dependent, differential AOX levels during early germination impact predictability of plant resilient performance (75, 157). **Figure 1B.1: Transcriptome profile of SARS-CoV-2 infected human lung adenocarcinoma cells (A549, MOI 0.2) 24 hours post infection (hpi)** **Results:** SARS-CoV-2 infection stimulated the immunological system, presented here by interferon regulator factor IRF9 (significant), and transcript factor NF-KB-RELA (112%, non-significant), although multiplicity infection rates were low (MOI 0.2) and *ACE2* and *TMPRSS2* could not be identified in A549 cells. Down-regulation of caspase initiator gene transcript levels, stable levels of caspase executor genes and up-regulated levels of *Bcl-xL* (all non-significant) are conform with arrested apoptosis activity in the host cell. Down-regulated *SNRK* (non-significant), unchanged transcript levels of *mTOR* and reduced *E2F1* cell cycle activator (non-significant) point together also to arrested cell cycle activity. This coincided with down-regulated *GAPDH* (significant), *Pyruvate Kinase* (non-significant) and *G6PDH* (significant) as well as *mt-MDH2* (non-significant). In this situation, *LDH* transcript level was found equal to control. However, transcript levels for *SOD2* as a biomarker for oxidative stress regulation was slightly increased (non-significantly) and for *GPX* as well as *GSR* down-regulated, whereas *SOD1* and *Catalase* showed control level. *ADH5*/*GSNOR* kept control level, but *NOS1* was slightly down-regulated (non-significant). This together with significantly down-regulated tubulin indicates the start of adaptive complex signaling and induced structural host cell reorganization. *IRF9* demonstrated early response of the immune system, which might qualify as functional marker candidate. *IRF3* remained at mock control level. *ACE2* (*Angiotensin-converting enzyme 2*), *TMPRSS2* (*transmembrane protease serine 2*) gene expression was not detected in the analysis and genes were not denoted in the Supplementary table S6. **Figure 1B.2: Transcript profiles of SARS-CoV-2 infected primary human bronchial epithelial cells (NHBE, MOI 2.0) and respiratory syncytial virus (RSV) - infected A549 cells (MOI 15) at 24hpi** **Results:** In NHBE cells, *ACE2* and *TMPRSS2* were identified. While the transcript level of *ACE2* remained unchanged, a significant decrease for ACE2-priming molecule TMPRSS2 was observed. Nevertheless, SARS-CoV-2 infection stimulated again the immunological system, presented here by *IRF9*, and *NF-KB1* (both significant). However, in this context different from **Figure 1B.1**, an equally strong increase in *SOD2* transcript level compared to *IRF9* was observed along with slightly up-regulated *SOD1*. This was accompanied by down-regulated levels of catalase, *GPX* and to a higher extent *GSR* (all non-significant) (Supplementary Table S7). Combining these last results, they signal changed oxidative stress level and complex fine-tuning activities. *NOS1* transcript level seems to be unchanged regarding the control. However, *ADH5*/*GSNOR* was down-regulated to 34% (significant), signaling a change in NO homeostasis. Increased *mTOR* transcript level and at the same time down-regulated *SNRK* level (both non-significant) coincide with significant *E2F1* transcript level increase, which goes along with a similarly strong transcript level increase of *Tubulin* (significant) and also of *Actin* (non-significant). Overall, this points to rapidly induced cell cycle activity. This picture is supported by an increase in *LDH* transcript level (significant) linked to an increase in glycolysis enzyme transcripts from *GAPDH* (significant) onwards and also *mt-MDH2* (non-significant), whereas *ct-MDH1* transcripts were reduced (non-significant). *G6PDH* transcript level was unchanged. The transcript level of anti-apoptotic *Bcl-xL* is significantly up-regulated, but also caspase initiator and caspase executor transcription were up-regulated though to an obviously lesser extent (both non-significant). In comparison, RSV-infected A549 cells responded under the applied experimental conditions strikingly similar in relation to the most pronounced host cell responses due to increased transcript levels of *mt-SOD2* and *IRF9* (number of replicates for transcripts was in part less for this experimental system, which can be responsible for missing significances). In both variants, *IRF3* remained basal. However, the overall response was differential and varying transcript level changes were also indicated for *NF-KB-RELA* and *NF-KB1*. Anti-oxidative enzyme transcript levels were more reduced apart from higher transcript accumulation for *GSR*. Also, *NOS1* transcripts were in this case down-regulated to 72.5% (non-significant). *ADH5*/*GSNOR* was reduced to a similar level (28%) as seen for SARS-CoV-2 infected NHBE cells. Although *LDH* transcripts were up-regulated to a similar extent as observed for SARS-CoV-2-infected NHBE cells, and again linked to up-regulation of the glycolysis pathway, the overall response was different. *GAPDH* transcript level was down-regulated and in this case, *Hexokinase* and *PFK-M* were up-regulated and enolase (significant) and pyruvate kinase showed higher transcript levels and *G6PDH* was down-regulated. A striking difference comes by the observation that *SNRK* was more strongly down-regulated, but *mTOR* was not up-regulated and *E2F1* did also not show up-regulation. This together linked to contrasting down-regulation of transcript levels for tubulin and obviously unchanged actin transcription. Collectively, it indicates that in this system at the given time point (24hpi), cell reprogramming was taking place, but cell cycle progress was not stimulated. No sign of cell death up- or down-regulation can be recognized. The transcript level of anti-apoptotic *Bcl-xL* is again significantly up-regulated, caspase initiator level remained basal and caspase executor transcription was down-regulated (non-significant). **Figure 1B.3: Transcriptome profiles of MERS-CoV – infected (MOI 4.0) human lung epithelial cancer cells (Calu-3) at 6 and 24hpi** **Results:** At 6hpi, *SOD2* transcript level was strongly up-regulated to 148% (significant) along with catalase (non-significant). *SOD1*, *GPX* and *GSR* were significantly down-regulated. *NOS1* seemed to be slightly up-regulated (non-significant) at slightly, but significantly reduced level for *ADH5*/*GSNOR*. However, at 24hpi, all transcript levels of oxidative stress-related enzymes and *NOS1* as well as *ADH5*/*GSNOR* were strongly down-regulated (all significant), which indicates drastic change in overall cell program. *SOD2* and *NOS1* were reduced to similar levels of around 40% of the control. *IRF9* transcript level was highly increased at 6hpi to about the same degree as *SOD2* (159% though non-significant), *IRF3* was significantly decreased along with a smaller increase in *NF-KB1* (non-significant). While *IRF9* was then further up-regulated at 24hpi to about 240% (significant), *IRF3* was further decreased along with *NF-KB1*, *NF-KB-RELA* (all significant) (Supplementary Table S8). At 6hpi a significant increase in *SNRK* (220% against the mock control) was the most enhanced signal among all components of the profile, which indicates strong early energy depletion crisis. However, at 6hpi *mTOR* transcripts were also significantly increased (to around 118%) along with a similar increase in *E2F1* (123%, significant), which signals rapid efforts for cell cycle progression despite significantly down-regulated, already low transcript levels for tubulin and actin. At that time, *LDH* remained basal and all transcript levels from glycolysis pathway, *G6PDH* and *MDH* were already reduced. *GAPDH* transcript level was most affected from glycolysis, but to a similar extent as *G6PDH*. At 24hpi, *SNRK* showed still increased transcript levels (143%; significant), but *mTOR* levels significantly decreased to around 70% of the control and all metabolism-related components were suppressed to levels below 50%. Down-regulation of *mTOR* at 24hpi associated with stronger down-regulation of all genes from glycolysis, most pronounced for *GAPDH*, *G6PDH* and both *MDH*s from cytoplasm and mitochondria. This was then also linked to strongly reduced *LDH* transcript level to 10% (significant) and more drastically reduced levels for tubulin (2%) and actin (1%). The anti-apoptotic gene *Bcl-xL* was down-regulated (non-significant) already at 6hpi and *Caspase* executor transcript level strongly up-regulated (significant) to 152%, which together signaled strong initial cell efforts to initiate apoptosis/cell death. However, at 6hpi caspase initiator was already down-regulated (significant) and at 24hpi all three cell death-related genes showed significantly reduced transcript levels. Thus, these combined results indicate that cell cycle had initially been rapidly progressed, but cell division and proliferation were not induced and apoptosis had been rapidly induced, but was suppressed with time. Nothing indicates at that time that energy-dependent metabolic reprogramming was substantially induced. **Figure 1B.4: Transcriptome profiles of MERS-CoV (a) and SARS-CoV (b) infected human fetal lung fibroblast cells (MRC5) by MOI 0.1/3.0 at 24hpi and 48hpi** **Results:** For MRC5 cells, ACE2 was not identified. MERS-CoV-infection observed at 24hpi and 48hpi (**Figure 1B.4a)** showed for all profile components down-regulation at different degrees of significance (see letters in figure). This included also *SOD2* and *IRF9* (both with significant reduction at 48hpi and low MOI). *NOS1* seemed to be slightly less affected (39.5%, significant) than mt-SOD2 (29%). However, *ADH5*/*GSNOR* levels are significantly reduced to 10%. At 24hpi, low MOI (0.1) showed within each component always highest transcript levels by comparing MOI and infection times. In general, at higher MOI the effect of time seemed to be reduced. *SNRK* responded exceptionally among all components. Transcript levels stayed comparatively more stable across all variants and had not been significantly reduced to control level at any time. *mTOR* showed a small reduction to the control at low MOI at 24hpi (non-significant), but transcripts were significantly reduced by time and also at higher MOI at 24hpi. *E2F1* showed a parallel pattern to *mTOR*, but was at 48hpi and low MOI more drastically reduced (25% to control) (significant) than *mTOR* (65% to control) (significant). Again, *GAPDH* was the most affected enzyme from the glycolysis path at both MOI (10.5% and 37.5%) and *G6PDH* was decreased to slightly lower extent (8% and 30%) (significant at 48hpi for both, low MOI). *Tubulin* and *Actin* revealed similarly drastic and significant reduction in transcript levels at low MOI 48hpi (to 12% and 10%). *LDH* transcript levels decreased with time at both MOI (significant for 48hpi, low MOI) (Supplementary Table S9). In SARS-CoV-infected MRC5 cells (**Figure 1B.4b**), we found from transcriptional ORFs much lower virus replication for SARS-CoV than for MERS-CoV (data not shown). This might have contributed to only moderate broad down-regulation in comparison to MERS-CoV infection. Under these conditions, we see in SARS-CoV infected cells up-regulated SOD2 at 48hpi at both MOI and differential down-regulation of most of the anti-oxidant components (non-significant). *NOS1* was downregulated with time and MOI (significant), but 48hpi at high MOI transcript levels were increased again to basal. Also *ADH5*/*GSNOR* transcript levels were down-regulated at higher MOI, but, in general, demonstrated slightly increased levels at 48hpi (non-significant). *IRF9* was slightly up-regulated (non-significant) only at 24hpi and higher MOI. *SNRK* was consistently up-regulated (113 - 126%, non-significant). To the contrary, initial *mTOR* values above the control were down-regulated with time at high MOI. Further, strong down-regulation was observed in mean values for *E2F1* at both MOI levels (non-significant). Tubulin and actin transcript levels also tended to decrease with time at both MOI levels. Though all these observations were separately not significant, together they suggest early energy depletion and suppression of cell cycle progression or cell proliferation at 48hpi. In agreement with this observation, *LDH* transcripts seemed to be unchanged to the control (no significant differences) (Supplementary Table S10). Nevertheless, a tendency of enhanced transcription of *LDH* was observed at 48hpi at both MOI, which went along with increased levels for enzymes related to glycolysis. Together, these observations indicate increased energy-dependent metabolic reprogramming with time. The anti-apoptosis gene *Bcl-xL* tends to be reduced with time. Both caspases remained at control level at low MOI, but both indicate a tendency for increasing transcription levels with time, which is more obvious, though non-significant, for the executing caspase (138%). Thus, it seems that apoptosis/cell death was stimulated by time.

### Bacterial endophyte and mycorrhiza treatments of *D. carota* L. seeds

Seeds of two carrot varieties, Kuroda and Early Nantes, were used for the experiments. At first, they were pretreated to remove native endophytes by an established protocol (75). Briefly, stock solutions of 1 mg ml^-1^ tetracycline (antibiotic) and miconazole nitrate (antifungal) were prepared and 5 ml of each solution was transferred onto a Petri plate. 200 mg of carrot seeds were weighed and added into the plate containing antibiotic and antifungal solutions and kept under constant shaking (100 rpm) for 6 hrs. Thereafter, seeds were washed with sterile deionized water for three times and kept under laminar air flow until they were completely dried. To confirm the effectiveness of the treatment, few seeds were collected randomly and cut into small pieces. These pieces were placed on nutrient agar/potato dextrose agar (Himedia, India) and incubated at 35 ^°^C/28^0^ C for overnight (bacteria) to 2 days (fungi). Absence of bacterial/fungal growth indicated that seeds were free from native endophytes.

For bacterial endophyte treatment, Single colonies of native bacterial endophytes (EN1, EN2, EN3) were inoculated in 100 ml of autoclaved nutrient broth (HiMedia, India) and incubated at 35 ^°^C overnight. Overnight grown bacterial cultures were centrifuged for 5 min at 4000 rpm to collect the pellet. The pellet is diluted with sterile deionized water until the optical density (O.D) reached to 0.2 (2 x 10^8^ cells ml^-1^). The endophyte-free seeds were immersed in 20 ml of each bacterial culture and kept under shaking at 100 rpm for 2 h. Then, seeds were dried under laminar air flow to remove excess moisture.

For arbuscular mycorrhizal fungi (AMF) treatments, 90 days old root organ culture (ROC; stock culture) of *Rhizophagus irregularis* and *Rhizophagus proliferus* was obtained from the Centre for Mycorrhizal Culture Collection (CMCC), TERI, India. The stock cultures of AMF1 and AMF2 were harvested at 25°C in 100 ml sodium citrate buffer using a shaker (Kuhner Shaker, Basel, Switzerland) (77). After deionization, buffer containing harvested spores (without roots) was sieved through a 325 British Standard (300 μm) Sieve (BSS) (Fritsch, Idar-Oberstein, Germany). Spores retained on the sieve mesh were washed with sterile distilled water (twice). Finally, the spores were collected in 20 ml of sterile distilled water (all steps were performed under aseptic conditions) (78).

For experimental set-up, autoclaved germination papers (Glasil Scientific Industries, New Delhi, India) were placed onto Petri dishes and moisturized with 5 and 10mM SHAM. Forty bacterial endophyte-treated seeds were placed onto dishes using sterile forceps. In the case of the AMF1 and AMF2 treatments, 40 endophyte free seeds were placed onto the dishes and 5 spores *of R. irregularis* were inoculated per seed. All treatments were setup with three replicates. Triplicates of controls were maintained by inoculating seeds in sterile distilled water. After inoculation, seeds were kept in darkness for 40 h and then transferred to light conditions. The plates were incubated in a culture room at 22-25°C with a 16h/8h photoperiod. Germinated seeds were counted and data were recorded at 24, 30, 48, 72, 96 and 120 HAI. Seed germination was identified by radicle emergence.

### Statistical analyses

Normality and homogeneity of variances from the samples were tested with Shapiro-Wilk test and Bartlett or Levene tests, respectively. If data were parametric, Student’s t-test (two populations) or one-way ANOVA were used. ANOVA was followed by Tuckeýs post hoc test when a significant effect of the tested factor was detected. When pre-requisites for parametric analyses were not kept, variable transformations were made (ln(x) and sqrt(x)) to perform parametric analyses, or non-parametric tests were used. These last tests were performed only for ADH levels in auxin-induced seeds (Wilcoxon test, **Fig. 1B**) and transcript levels obtained after viral infections of MRC5 cells (Krukal-Wallis test, **Fig. 1B.4**). After ANOVA, post hoc analyses (Tukey’s) were performed. Statistical packages used were InfoStat v. 2018 and R v. 4.0.2.

We highlight that we interpret our data as ‘real’ observations under the employed conditions involving only small samples, which certainly provide insights that cannot get relevance or not relevance by using significance calculation. Nevertheless, we applied significance calculations at usual p-values for biological research as an aid to appropriately focus our insights. This is the reason for describing our observations in figure legends by reporting the attributes of the data as either significant or non-significant only as additional information in parenthesis. Readers are encouraged to making themselves familiar with the current paradigm change related to the usage of statistical significance (79–83).

## Results & Discussion

Following the perspective of Arnholdt-Schmitt et al. (21, in press), our principle goal of this study on coronaviruses with special reference to SARS-CoV-2 was the validation of the hypothesis that transcript level profiles of justified target genes established from *in vitro* somatic embryogenesis (SE) induction in plants compared to virus-induced profiles can identify differential virus signatures that link to harmful reprogramming.

Thus, we first established a reference transcript profile from a plant model experimental system and subsequently compared this reference profile step-by-step along the outlined five characteristics with transcriptome profiles of corona virus-infected cells. As a consequence, we were able to extract targets promising for developing a strategy to avoid virus spread already in primary infected cells.

In **Figure 1**, we present functional transcriptome profiles related to reprogramming under SE-inducing conditions in the model species *Arabidopsis thaliana* (**Figure 1.A.1**) and from corona virus-induced human cell lines (**Figure 1.B.1-4**). Results of detailed observations and complementing wetlab research results are highlighted in the subfigures and in **Figure legend**. In the main text, we focus on the principle line of this research.

**In summary,** we found (**A**) that the reference plant transcript level profile marked early *de novo* cell programming by five salient characteristics as follows:

(**1**) Modified complex oxidative stress signaling pattern and a special role for increased superoxide dismutase (SOD) as stress indicator
(**2**) Decreased transcript levels of NO-producing NR
(**3**) Signals that indicate arrested cell cycles at reduced alpha-tubulin transcript levels
(**4**) Transient increase in aerobic fermentation connected to enhanced glycolysis
(**5**) Activation of the cell death-regulating system without cell death down-regulation

By comparing this reference profile step-by-step along these outlined five characteristics with transcriptome profiles of corona virus-infected cells, we **(B)** noted the following observations:

**(for 1)** increased *SOD2* transcript levels marked also early corona viral infection in cultured human cells (MERS-CoV, SARS-CoV and SARS-CoV-2). However, considering the pattern of anti-oxidant enzymes, virus-infected cells responded with differential complex patterning depending on infection pressure, hours post infection, virus type and host cell type;

**(for 2)** in MERS-CoV infected calu-3 cells at MOI 4.0, up-regulation of *NOS3* transcript level was observed already at 6hpi, which strongly increased at 24hpi. However, in SARS-CoV-2 infected NHBE cells at MOI 2.0, no transcription of *NOS* genes was observed;

**(for 3)** as in the reference profile, SARS-CoV-2 infected A459 cells at low MOI (0.2) signaled arrested cell cycle at significantly reduced alpha-tubulin transcript levels. However, SARS-CoV-2 infected NHBE cells at 10 times higher MOI (2.0) signaled cell cycle progression combined with significant increase in tubulin transcript levels. Also, MERS-CoV-infected calu-3 cells at high MOI (4.0) signaled cell cycle progression early at 6hpi, but in the presence of already significantly down-regulated tubulin transcripts. At 24hpi, arrested cell cycles were indicated, which was then combined with more drastic down-regulation of tubulin transcript levels. In agreement with the latter result, in MERS-CoV- and SARS-CoV-infected MRC5 cells at 24hpi and 48hpi, no sign of cell cycle progression or proliferation was found, which was independent on lower (0.1) or higher (3.0) MOI. Cell cycle arrest-signaling was combined with down-regulated tubulin transcript levels as also seen in the reference profile;

**(for 4)** SARS-CoV-2 infected NHBE cells (MOI 2.0) signaled significant increase in aerobic fermentation through increased *LDH* transcript levels, which was linked to enhanced transcript levels for enzymes associated with glycolysis. In MERS-CoV-infected calu-3 cells (MOI 4.0) at 6hpi, *LDH* levels were found to be basal. This was linked to reduced transcript levels for glycolysis. At 24hpi, transcript levels of *LDH* and glycolysis related genes were drastically reduced

**(for 5)** SARS-CoV-2 infected A549 cells at low MOI (0.2) indicated cell death arrest at 24hpi. NHBE cells that were infected at 2.0 MOI with SARS-CoV-2, signaled at 24hpi activation of the cell death-regulating system. RSV-infected A549 cells at 15 MOI indicated at 24 hpi cell death arrest. MERS-CoV infected calu-3 cells (MOI: 4.0) indicated strong induction of the cell death regulatory system at 6hpi and suppression of the system at 24hpi. Also, in MERS-CoV infected MRC5 cells, suppression for the cell death-regulating system was observed at 24hpi and at 48hpi at MOI 0.1 and 3.0. SARS-CoV infected MRC5 cells under the same experimental conditions, showed much lower replication rates than MERS-CoV infection (transcriptional ORF reads, not shown). It might be due to this context that initial activation of the cell death regulatory system is indicated at 24hpi also in SARS-CoV-2 infected cells, but that this initial activation was still followed at 48hpi by a trend for increasing caspase (initiator and executor) transcript levels.

**As a consequence from our observations described under A and B**, we **(C)** could extract as main result the following targets promising for developing a strategy to avoid early virus spread:

1) Balanced pattern of oxidative stress pattern
2) Decreased NO production
3) Avoidance of cell cycle progression by disconnecting increased aerobic fermentation from energy canalization to alpha-tubulin-based cell restructuration early during viral infection
4) Avoidance of prolonged cell death promotion
5) Considering the unique holobiont nature of individuals in firstly virus-infected cells

### Taken together, this complex approach resulted in arriving at two principle conclusions

**-** Comparing *in vitro* coronavirus-induced transcriptome profiles with plant cell transcript profiles during *de novo* programming as a reference enabled to identify main characteristics for early SARS-CoV-2-induced transcript changes. Collectively, they indicate one *m*ajor *c*omplex *t*rait for *e*arly *de novo* programming, named here as ‘CoV-MAC-TED’: unbalanced ROS/RNS levels connected to increased aerobic fermentation that links to alpha-tubulin-based cell restructuration and cell cycle progression.

- In plant systems, it could be shown that the extent of aerobic fermentation induced during *de novo* programming that linked to the initiation of embryonic or non-embryonic cell proliferation was regulated by interacting sucrose- and AOX-levels. Early up-regulation of alcoholic and lactic aerobic fermentation was connected to higher glycolysis and oxidative stress levels. This was associated with increased *AOX* transcript accumulation. Furthermore, our results suggested that a mutant’s capacity for more efficient reprogramming compared to wild type (WT) was linked to the capacity of limiting aerobic fermentation, which associated positively to *AOX* transcription levels.

These results are in good agreement with several studies, which showed the occurrence of ROS/RNS disturbances during early viral infection, a potential role of hijacked aerobic fermentation for virus replication, involvement of cytoskeleton during viral infection and virus-induced cell cycle modulation (see in background). The findings further pinpoint to the beneficial role of AOX for plant resilience that is related to both ROS/RNS equilibration and redox homeostasis, able to avoid acidification and excess of toxic ethanol, and regulating at the same time adaptive energy supply for growth performance (21, in press, 84, preprint). Interestingly, Ito et al. (85) discovered in *Arum* that temperature-dependent switching between critical *AOX* polymorphisms in the binding site for AOX-pyruvate can determine energy-related metabolic regulation, which in turn results in plant performance. This striking finding underlines again the multifunctional role of AOX that includes besides ROS/RNS equilibration adjusting energy metabolism at the threshold between mitochondrial respiration and aerobic fermentation. *AOX* polymorphisms have been identified and widely explored as genetic trait (86–96), besides being epigenetic (97) and developmental manifestation (98, 99). Also, polymorphisms in neighboring regions of conserved functional sites had been shown to discriminate AOX isoenzymes (100).

Furthermore, it was found that *AOX* polymorphisms could be used to distinguish individual plants from the same species (101). Symbiotic AMF revealed substantial *AOX* gene polymorphism within and between spores (102), an observation which awaits to be functionally explored in relation to adaptive plant holobiont performance (103–107).

Our results demonstrated that endophytes can interfere with the redox biology of the host system related to the initiation of cell proliferation (**Fig.1A.4**). Plant-mycorrhizal fungi interaction is suggested to involve *AOX* from both symbiotic partners (102–104, 106). Consequently, the holobiont nature of primary virus-infected cells should be considered as influencer on the impact of early virus infections on cell redox status. We suppose that endophyte interaction is important for program initiation/realization rather than for early program induction (75, 108). This view is in agreement with Visser-Tenyenhuis et al. (109), who observed enhanced SE by bacterial co-cultivation, although bacteria *per se* could not induce SE. Bharadwaj et al. (84, preprint) extended here reported results from endophyte effects during germination and could show that sucrose critically affected early endophyte impacts on the initiation of cell division growth dependent on the quality of microbiota. It was concluded that endophytes and symbiotic fungi can buffer negative effects of excess in sucrose during early reprogramming. In this way, microbiota and AOX are supposed to interact in support of equilibrated rapid adaptation to sugar-transmitted inner and outer environment signaling (see concept in 84, preprint).

The sophisticated role of *AOX* in plants had developed along evolution in the context of complex holobiont systems. Currently, AOX is developed as a tool for understanding the functionality of its beneficial mechanisms also in mammals that naturally do not enclose AOX (see references in 21, in press). It showed good integration into normal physiology when constitutively overexpressed in animals. However, mammals did not evolve AOX genes as an integral part in their complex metabolic and multi-organism networks. Thus, whether AOX could be useful in therapy as proposed for mitochondrial respiratory deficiency diseases (110–115)) will, in our view, crucially depend on its adaptive (positive and negative) regulation, which includes also interaction with endophytes. In this context, it is also pertinent to mention that recently an AOX-degrading protein has been discovered that might be added as a tool in potential AOX-based therapies (85).

In search for similarities between the beneficial role of AOX in plants in relation to adaptive oxidative stress level equilibration relevant for virus tolerance and that of natural agents in human cells, melatonin seems to be a strong candidate. Melatonin is a natural hormone in humans, which has also been recognized as a phytohormone (116). It is produced in most organs and cells (70, 116–119), including also human salivary gland cells (120). Melatonin is known to possess anti-oxidant properties and shows high fluctuation in its cellular concentration. In plants, melatonin seems not to enhance *AOX* transcription (116) as it has been shown for auxin (75). In turn melatonin interacts with other enzymes involved in ROS/RNS balancing and was suggested to be in plants mainly involved in biotic stress defense rather than in growth regulation (116). Further, melatonin was reported to enhance the induction of adventitious roots through interaction with auxin-mediated signaling, which suggests a role for melatonin also in early reprogramming (121). All evidence put together, it could be suggested that melatonin might substitute early anti-oxidant function of AOX during reprogramming (75, 99, 122). By looking at the transcription of melatonin synthesis-related genes *ASMT* (*N*-acetylserotonin O-methyltransferase) and *NAT* (serotonin *N*-acetyltransferase) in WT *A.thaliana* our model system showed that auxin-induced *AOX* transcript level changes, we found up-regulated *ASMT*/*NAT* transcript levels 55 hours after 2,4-D treatment though at a low level (not shown). This preliminary observation might lead to future interest and encourages further studies. Currently, the molecular-physiological and clinical relevance of melatonin is found to be a hot topic. It was highlighted to show important functionality in physiology, pathophysiology and chronobiology. Since long melatonin was also recognized as a beneficial agent for managing viral infections (123). However, despite its widespread use as a drug for many other purposes, its functional role and application still needs stronger confirmation through extensive biochemical and clinical research (124–126). Apart from acting as an anti-oxidant, melatonin demonstrates anti-inflammatory activity and immune-enhancing features besides interacting with ACE2 (123, 126–128). Recently, the proposed rationale for employing melatonin as a potential anti-viral agent in general and anti-SARS-CoV-2 agent in particular has been indicated by Zhang et al. (126). Extensive efforts are being made to identify drugs for treating SARS-CoV-2 infections via a network-based drug repurposing (126). By adopting this methodology, melatonin has been identified as a promising drug, which can be administered either alone or in combination with immune-suppressant agents. In our studies, we focused additionally on melatonin-synthesis related genes to work-out its drug potentials. Results of the present study showed enhanced *ASMT* transcript accumulation at low level in SARS-CoV-2-, RSV-, MERS-CoV- and SARS-CoV-infected human cells of various origins (**Fig. 2A** and **2B**). Related to MERS-CoV-infected MRC5 cells, it can be seen that *ASMT* transcript levels were increasing dependent on MOI level and infection time (**Fig. 2B**). However, we did not identify *ASMT* transcripts in MERS-CoV-infected calu3 cells (MOI 4.0). Nevertheless, these observations might encourage further research on the significance of melatonin actions during early viral infection in the primarily affected nose and mouth cells.

**Figure 2:**
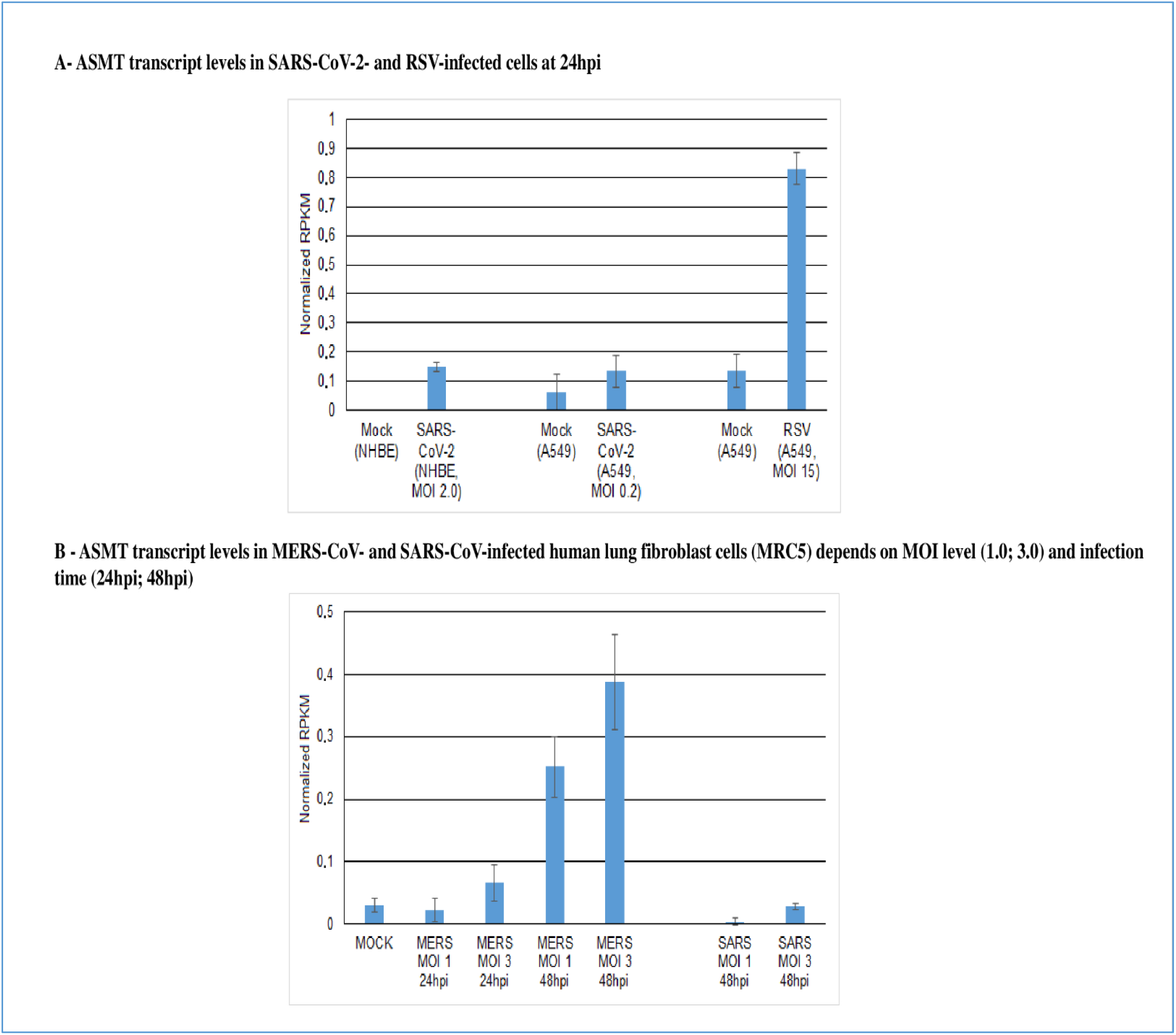
Expression of *ASMT* (acetylserotonin O-methyltransferase) gene involved in melatonin biosynthesis in virus-infected human cells.

Another interesting approach in agreement with the focus of our study is the potential anti-viral use of ergothioneine (ET) (129). ET is a natural amino acid that enters via diet into the human body and can be found in all animal cells. It has multifunctional characteristics of interest for anti-viral functionality that includes flexible regulation of redox biology. ET makes an attractive option for its use as an alternative to AOX functionality in plants essentially due to its highly plastic distribution based on an appropriate adaptive transporter system (OCTN1) and its ubiquitous availability. However, in order to be functional during the early phase in mouth and nose cells ET needs to have direct effect on avoiding virus-induced reprogramming that connects to initiating viral replication, which awaits future research (129).

In plants, impressive success has been achieved in confronting viral infections by applying TOR inhibitors or TOR silencing (reviewed in 55). Avoiding cell cycle progression in primary virus-threatened nose and mouth cells could be critical during the early phase of virus entry. Drugs that target mTOR and tubulin are available (50, 128) and they might be repurposed and adapted to early treatment of primary corona virus-threatened cells in mouth and nose. However, prolonged suppression of cell renewal in nose and mouth could be fatal due to collateral bacterial or other infections or contaminations. Consequently, pulsed treatment strategies could be considered to avoid virus-induced cell progression that benefits virus replication, but at the same time allowing intermittent relaxation from cell cycle inhibition for cell renewal. This type of treatment might enable ‘growing-out of danger’ of host mouth/nose cells. This is a general strategy followed in plants *in vivo* and in plant biotechnology systems in order to get rid of virus spread. Comparably, pulsed treatment strategy might contribute to stimulation of cell proliferation in a way that virus replication cannot follow in the same velocity as healthy new cells are emerging.

In plant biotechnology, it is well established that phytohormones and their relative intracellular concentrations are essentially involved, either directly or indirectly, in program decisions about growth and differentiation. However, the impact of phytohormones depends on available bioenergy as the critical frame for program realization. Phytohormones are known to be critical for *de novo* membrane-based cell restructuring (‘innovation’) during quiescent cell reprogramming, whereas sucrose is required for structural organization (‘performance’) (130). In human cells, the entry receptor of SARS-CoV-2, i.e. ACE2, is also known to function as a negative regulator of the hormonal renin-angiotensin system (RAS). RAS plays an important role in regulating cellular growth, proliferation, differentiation, and apoptosis besides extracellular matrix remodeling and inflammation (131). The receptor has been shown to protect lung from injury (132). However, for SARS-CoV infection and the Spike protein it had been shown that they reduce *ACE2* expression. Spike injection into mice worsened acute lung failure *in vivo*, which could be attenuated by blocking the renin-angiotensin path (133). Actual studies are being focused to enable efficient anti-SARS-CoV-2 treatment via competing soluble ACE2 drugs (hrsACE2) (132). We are in favor of this approach and suggest extending studies on hrsACE2 by considering the early indicated metabolic effects, which are presented as complex trait ‘CoV-MAC-TED’.

Rodríguez et al. (134) showed that differential action of phytohormones on the induction of new program (adventitious organogenesis) was epigenetically regulated. RAS action has also been linked to changes in the epigenome (135). During auxin-induced *de novo* cell programming, genome-wide hyper-methylation is known to take place (firstly reported for SE by 136). Gross hyper-methylation combined with genome-wide reduction in the amount of repetitive DNA linked to the occurrence of small DNA fragments had systematically been observed before cell cycle progression started (137). Therefore, we would like to suggest exploring the effect of soluble ACE2 drugs also on the level of epigenetic events that occur early during reprogramming before the cell cycle activation. Genome-wide repetitive DNA reduction linked to an increase in small DNA-fragments was also reported to happen before actual cell cycle start as a determinant for cell proliferation (137–139) and is supposed to be another powerful candidate linked to the induction of cell proliferation.

## Conclusion

This is the first research to start validating hypothesis and wider perspective published by a non-institutional, voluntary competence focus under the title ‘From plant survival under severe stress to anti-viral human defense – a perspective that calls for common efforts’ (21, in press).

We identified a corona virus-induced complex trait named ‘CoV-MAC-TED’ (*co*rona*v*irus *ma*jor *c*omplex *t*rait for *e*arly *de novo* programming). It is well established during first few hours of viral infection and consists of unbalanced ROS/RNS levels connected to increased aerobic fermentation that links to alpha-tubulin-based cell restructuration and cell cycle progression.

Primary infected nose and mouth cells become rapidly ‘super-spreaders’. This can have serious consequences for neighboring cells, the infected organism and for the environment.

Virus structures as non-living particles would be harmless, if the entry cells could ignore their presence. It is the proper host cell-reaction that is required and ‘abused’ for virus replication. Consequently, as long as we cannot avoid virus structures from establishing contact ‘with us’, we need to support the entry cells to become ignorant. If possible, this should happen proactively or, at least very rapidly and much before any symptoms of viral infection appears.

Considering our results, the first fingerprint of viral infection is complex and needs combined strategies that rigorously target the components of ‘CoV-MAC-TED’ in primary infected nose and mouth cells.

Thus, it is the need of the hour to develop efficient strategies to early equilibrate or rebalance ROS/RNS levels and to control cell progression in nose and mouth cells. We have discussed some promising approaches that are in agreement with our observations.

The possibilities of using controlled pH to support these strategies by early testing and treatment, should, in our view, be explored with priority. Cell reprogramming and its relation also to cell death (140), changes in anti-oxidant activity also related to AOX (141–143), aerobic fermentation (144) and viral infections (144–148) are all influenced or indicated by extracellular pH. Extracellular pH might also by its interaction with intracellular pH influence complex tasting, which includes not only sour tasting (149, 150). The relation between smell and pH is, to our knowledge, not sufficiently explored yet. However, SARS-CoV-2 entry proteins were found to be expressed in olfactory epithelial cells (151).

The prominent involvement of stress-related redox biology (see also 152) and importance of equilibration at very early steps of viral infection, calls pragmatically for respecting human individuality in its actual complex living and health situation at all processes of disease and social handlings. This challenges sampling procedures and treatment performance. For example, tests developed for diagnosis that require merely soft mouth and/or nose rinsing for sampling by the proper client in a quiet place are surely more beneficial compared to uncomfortable and thus, stressful sampling by medical assistance in an open space under time pressure. It also calls for appropriate social communication in order to support transparency and constructive information and avoid unnecessary creation of panic (153). These aspects should not be ignored and seen as meaningless ‘luxury’, but rather considered as being essential part of an overall strategy that aims primarily to seriously promote healthiness on the basis of current scientific knowledge and, at the same time, to prevent unbalanced stress factor development which is known to worsen health.

It was the driving force of our own cultural development, which resulted in the vicious cycle of non-sustainable growth and self-destruction. Settling of human populations required intensive and safe local food and feed production and at the same time allowed human populations to grow much more rapidly. This in turn increased human population’s need for safe (bio-)energy production per unit area. Science and technology development helped to fulfill this requirement and, thus, supported ‘unlimited growth’ at all levels. Plant and animal breeding for bioenergy-rich, quality-defined biomass are never ending efforts to permanently adapt domesticated organisms to changing natural and human-dominated world. Industrial agriculture is a modern history of innovative energy transformation processes. This overall phenomenon dedicated to growth, finally, led to dense human and domesticated animal populations within the restricted territory of Earth. It was followed by increased mobility and interaction between populations and accelerated the exchange of goods, but also caused rapid global spread of viruses and pathogens potentially harmful to plants, humans and other animals in the ecosystem. Major effects of human behavior on climate changes and potential interactions with virus spread are now in the focus of current awareness (154–156). To break this cycle of self-destruction in a non-violent, joy- and peaceful way and turn the rolling circle into a virtuous cycle of self-controlled sustainable growth and development, which includes also necessity for healthy aging, holistic education is needed. Holistic formation enables to recognize that sustainability is based on diversity as a stabilizing factor for robust natural and socio-economic ecosystems. Targeting these goals requires self-respect and a healthy balanced respect for ‘the other’ – inside and outside of us at all levels.

## Author Contributions

JHC coordinated and performed transcriptome analyses supported by KTL and SA and developed together with BAS the scientific approach, GM performed carrot SE lab analyses and helped together with BR in figure design and formal manuscript organization, BR together with SS and AA were responsible for lab studies on endophytes and AMF. SRK together with BR and GM analyzed the experimental data and helped BAS in overall *FunCROP* group coordination. BR wrote together with BAS part of the manuscript and prepared overall manuscript for submission. RS supervised lab work of GM and BR. CN by support of MO performed all statistical analyses and was strongly engaged to improve performance and data interpretation. KJG together with AK helped designing manuscript parts related to NO metabolism. All co-authors read, commented and discussed together or in smaller groups the overall manuscript and agreed on final submission.

## Acknowledgments

JHC is grateful to CNPq for the Researcher fellowship (CNPq grant 309795/2017-6). GM is grateful to UGC, India, for doctoral grant from BSR fellowship. GM, RS, and BAS acknowledge support for academic cooperation and researcher’s mobility by the India-Portugal Bilateral Cooperation Program (2013–2015), funded by “Fundação para a Ciência e Tecnologia” (FCT), Portugal, and the Department of Science and Technology (DST), India. BR and SS acknowledge the infrastructure and stay support provided by DBT-TDNBC-DEAKIN – Research Network Across continents for learning and innovation (DTD-RNA) for AMF related work at The Energy and Resources Institute, TERI, India. CN acknowledges the international scientific network BIOALI-CYTED, which contributed to establish FunCROP contacts. KTL is grateful to CNPq for the Doctoral fellowship. SA is grateful to CAPES for the Doctoral fellowship. KJG, MO and BAS acknowledge support by the India-Portugal Bilateral Cooperation Program ‘DST/INT/Portugal/P-03/2017’. MO Research is partially supported by National Funds through. FCT. Fundação para a Ciência e a Tecnologia, projects UIDB/04674/2020 (CIMA). BAS wants to thank Dr. Natascha Sommer for helpful discussions and comments on the manuscript during its development on the background of her experience as medical doctor in the group of Prof. Dr. Norbert Weissmann, Chair for ‘Molecular Mechanisms of Emphysema, Hypoxia and Lung Aging’ at the Universities of Giessen and Marburg Lung Center (UGMLC), Germany, and as investigator involved in mitochondrial redox biology also by help of transgenic AOX-mice. BAS acknowledges the strong engagement and clear view of Dr. Kim Berit Lewerenz-Kemper transmitted in support of our approach and final wider conclusions that is based not only on her overall life experience and view but also on her coronavirus-dominated experience as medical doctor in emergency service at the hospital ‘Universitätsklinikum Eppendorf’ in Hamburg, Germany. Furthermore, we want to appreciate encouragement and comments on the manuscript by Dr. Elisete Santos Macedo, Prof. Dr. Ashwani Kumar (emeritus at the University of Jaipur, India) and Dr. Nidhi Gupta, all are engaged plant scientists, educators, book writers and members of the FunCROP net and accompanied developing this approach from beginning. Finally, the authors thank Sebastian Schaffer, Nutritional Scientist with special knowledge in anti-oxidant food ingredients and their relevance in aging and as neuroprotective agents, who currently founded ACADELION Scientific and Medical Communications, for his carefully reading and helpful editing of the manuscript.

## Supplementary figures

**Supplementary figure S1:** Differences in transcript levels (normalized in RPKM values) of *AOX*, *ADH1*, *LDH* and *Enolase* in WT and clf/cwn-mutant between mock controls and between 55h auxin-treated seeds. **: highly significant (p < 0.01) differences

**Supplementary figure S2:** Differences in transcript levels (normalized in RPKM) of *AOX1* and *AOX2* between mock control and auxin-treated seeds of Arabidopsis thaliana (**: highly significant (p < 0.01))

## Supplementary Tables

**Supplementary Table S1**: Mean values ± SE corresponding to the **figure 1A.1**. Number of biological replicates: 3. *AOX* (a*lternative oxidase*), *ADH* (*alcohol dehydrogenase*), *LDH (lactate dehydrogenase)*, *Cyt-Fe-SOD1* (*Cytosolic-iron-Superoxide dismutase 1*), *mt-Mn-SOD1* (*mitochondrial manganese superoxide dismutase 1*), *CAT3* (*Catalase 3*), *GPX* (*Gluthatione peroxidase*), *Cyt-GSR1* (*Cytosolic-glutathione reductase 1*), *PAL* (*Phenyl alanine ammonia lyase*), *CHS* (*Chalcone synthase*), C3H (*p-coumarate 3-hdroxylase*)

**Supplementary Table S2**: Mean values ± SE corresponding to the **figure 1A.1**. Number of biological replicates: 3. *CAD* (*cinnamyl alcohol dehydrogenase*), *NR* (*nitrate reductase*), *ADH2* (*alcohol dehydrogenase 2*), *SNRK* (*sucrose non-fermenting related kinase*), *mTOR* (*mammalian target of rapamycin*), *E2F1* (Transcription factor E2F), *BAG* (*Bcl-2 associated gene*), *MC* (*Meta-caspase*).

**Supplementary Table S3**: Mean values ± SE of transcript abundance of Alternative oxidase (*AOX*) and Alcohol dehydrogenase (*ADH*) genes in *Arabidopsis* seedlings treated with auxin. The graphical representation of the values is given in **figure 1A.1**. Number of biological replicates: 3.

**Supplementary Table S4**: Mean values ± SE of transcript abundance of *BAG* (*Bcl-2 associated gene*) in *Arabidopsis* seedlings treated with auxin. The graphical representation of the values is given in **figure 1A.1**. Number of biological replicates: 3.

**Supplementary Table S5**: Mean values ± SE of transcript abundance of *MC* (*Meta-caspase*) gene in *Arabidopsis* seedlings treated with auxin. The graphical representation of the values is given in **figure 1A.1**. Number of biological replicates: 3.

**Supplementary Table S6**: Mean values ± SE of transcript abundance of genes in human lung adenocarcinoma cells infected with SARS-CoV-2. The graphical representation of the values is given in **figure 1B.1**: Number of biological replicates: 3. *LDH* (*lactate dehydrogenase*), SOD1 (*superoxide dismutase1*), SOD2 (*superoxide dismutase 2*), *GPX* (*glutathione peroxidase*), *GSR* (*glutathione reductase*), *ADH5* (*alcohol dehydrogenase 5*), *PFK* (*Phosphofructokinase*), *GAPDH* (*Glyceraldehyde-3-phosphate dehydrogenase*), *G6PDH* (*glucose-6-phosphate dehydrogenase*), Cyt-MDH1 (*Cytosolic malate dehydrogenase 1*), *mt-MDH2* (*mitochondrial malate dehydrogenase*), *SNRK* (*sucrose non-fermenting related kinase*), *mTOR* (*mammalian target of rapamycin*), E2F (Transcription factor E2F), *IRF9* (*interferon regulatory factor 9*), *IRF3* (*interferon regulatory factor 3*), *NFKB1* (*nuclear factor kappa B1*), *NFKB-RelA* (*nuclear factor Kappa B-RelA*), *Caspase In* (Initiator caspases), Caspase Ex (*Executioner caspases*), *Bcl-xL*, (*B-cell lymphoma-extra large*). *ACE2* (*Angiotensin-converting enzyme 2*), *TMPRSS2* (*transmembrane protease serine 2*) gene expression was not detected in the analysis and genes were not denoted in the table.

**Supplementary Table S7**: Mean values ± SE of transcript abundance of genes in primary human bronchial epithelial cells infected with SARS-CoV-2. The graphical representation of the values is given in **figure 1B.2**. Number of biological replicates: 2-3. *LDH* (*lactate dehydrogenase*), SOD1 (*superoxide dismutase1*), SOD2 (*superoxide dismutase 2*), *GPX* (*glutathione peroxidase*), *GSR* (*glutathione reductase*), *ADH5* (*alcohol dehydrogenase 5*), *PFK* (*Phosphofructokinase*), *GAPDH* (*Glyceraldehyde-3-phosphate dehydrogenase*), *G6PDH* (*glucose-6-phosphate dehydrogenase*), Cyt-MDH1 (*Cytosolic malate dehydrogenase 1*), *mt-MDH2* (*mitochondrial malate dehydrogenase*), *SNRK* (*sucrose non-fermenting related kinase*), *mTOR* (*mammalian target of rapamycin*), E2F (Transcription factor E2F), *IRF9* (*interferon regulatory factor 9*), *IRF3* (*interferon regulatory factor 3*), *NFKB1* (*nuclear factor kappa B1*), *NFKB-RelA* (*nuclear factor Kappa B-RelA*), *Caspase In* (Initiator caspases), Caspase Ex (*Executioner caspases*), *Bcl-xL*, (*B-cell lymphoma-extra large*) *ACE2* (*Angiotensin-converting enzyme 2*), *TMPRSS2* (*transmembrane protease serine 2*).

**Supplementary Table S8**: Mean values ± SE of transcript abundance of genes in primary human lung epithelial cancer cells infected with MERS-CoV. The graphical representation of the values is given in **figure 1B.3**. Number of biological replicates: 3. *LDH* (*lactate dehydrogenase*), SOD1 (*superoxide dismutase1*), SOD2 (*superoxide dismutase 2*), *GPX* (*glutathione peroxidase*), *GSR* (*glutathione reductase*), *NOS3* (*nitric oxide synthase*), *ADH5* (*alcohol dehydrogenase 5*), *PFK* (*Phosphofructokinase*), *GAPDH* (*Glyceraldehyde-3-phosphate dehydrogenase*), *G6PDH* (*glucose-6-phosphate dehydrogenase*), Cyt-MDH1 (*Cytosolic malate dehydrogenase 1*), *mt-MDH2* (*mitochondrial malate dehydrogenase*), *SNRK* (*sucrose non-fermenting related kinase*), *mTOR* (*mammalian target of rapamycin*), E2F (Transcription factor E2F), *IRF9* (*interferon regulatory factor 9*), *IRF3* (*interferon regulatory factor 3*), *NFKB1* (*nuclear factor kappa B1*), *NFKB-RelA* (*nuclear factor Kappa B-RelA*), *Caspase In* (Initiator caspases), Caspase Ex (*Executioner caspases*), *Bcl-xL*, (*B-cell lymphoma-extra large*).

**Supplementary Table S9**: Mean values ± SE of transcript abundance of genes in primary human fetal lung fibroblast cells infected with MERS-CoV. The graphical representation of the values is given in **figure 1B.4 a**. Number of biological replicates: 3. *LDH* (*lactate dehydrogenase*), SOD1 (*superoxide dismutase1*), SOD2 (*superoxide dismutase 2*), *GPX* (*glutathione peroxidase*), *GSR* (*glutathione reductase*), *ADH5* (*alcohol dehydrogenase 5*), *PFK* (*Phosphofructokinase*), *GAPDH* (*Glyceraldehyde-3-phosphate dehydrogenase*), *G6PDH* (*glucose-6-phosphate dehydrogenase*), Cyt-MDH1 (*Cytosolic malate dehydrogenase 1*), *mt-MDH2* (*mitochondrial malate dehydrogenase*), *SNRK* (*sucrose non-fermenting related kinase*), *mTOR* (*mammalian target of rapamycin*), E2F (Transcription factor E2F), *IRF9* (*interferon regulatory factor 9*), *IRF3* (*interferon regulatory factor 3*), *NFKB1* (*nuclear factor kappa B1*), *NFKB-RelA* (*nuclear factor Kappa B-RelA*), *Caspase In* (Initiator caspases), Caspase Ex (*Executioner caspases*), *Bcl-xL*, (*B-cell lymphoma-extra large*).

**Supplementary Table S10**: Mean values ± SE of transcript abundance of genes in primary human fetal lung fibroblast cells infected with SARS-CoV. The graphical representation of the values is given in **figure 1B.4 b**. Number of biological replicates: 2-3. LDH (lactate dehydrogenase), SOD1 (superoxide dismutase1), SOD2 (superoxide dismutase 2), GPX (glutathione peroxidase), GSR (glutathione reductase), ADH5 (alcohol dehydrogenase 5), PFK (Phosphofructokinase), GAPDH (Glyceraldehyde-3-phosphate dehydrogenase), G6PDH (glucose-6-phosphate dehydrogenase), Cyt-MDH1 (Cytosolic malate dehydrogenase 1), mt-MDH2 (mitochondrial malate dehydrogenase), SNRK (sucrose non-fermenting related kinase), mTOR (mammalian target of rapamycin), E2F (Transcription factor E2F), IRF9 (interferon regulatory factor 9), IRF3 (interferon regulatory factor 3), NFKB1 (nuclear factor kappa B1), NFKB-RelA (nuclear factor Kappa B-RelA), Caspase In (Initiator caspases), Caspase Ex (Executioner caspases), Bcl-xL, (B-cell lymphoma-extra large).

**Supplementary Table S11**: Bioproject details of RNA-seq experiments used to obtain the gene expression in *Arabidopsis thaliana* and Human cells

