## Supplementary Tables S1 to S11 for "ROS/RNS balancing, aerobic fermentation regulation and cell cycle control – a complex early trait (‘CoV-MAC-TED’) for combating SARS-CoV-2-induced cell reprogramming"

**Supplementary Table S1:** Mean values ± SE corresponding to the figure 1A. Number of biological replicates: 3. *AOX* (a*lternative oxidase*), *ADH* (*alcohol dehydrogenase*), *LDH (lactate dehydrogenase)*, *Cyt-Fe-SOD1* (*Cytosolic-iron-Superoxide dismutase 1*), *mt-Mn-SOD1* (*mitochondrial manganese superoxide dismutase 1*), *CAT3* (*Catalase 3*), *GPX* (*Gluthatione peroxidase*), *Cyt-GSR1* (*Cytosolic- glutathione reductase 1*), *PAL* (*Phenyl alanine ammonia lyase*), *CHS* (*Chalcone synthase*), C3H (*p-coumarate 3-hdroxylase*)

| **Treatment** | **Total *AOX*** | ***ADH1*** | ***LDH*** | ***ADH*** | **Enolase** | ***Cyt-Fe-SOD1*** | ***mt_Mn-SOD1*** | ***CAT3*** | ***GPX*** | ***Cyt-GSR1*** | ***PAL*** | ***CHS*** | ***C3H*** |
| --- | --- | --- | --- | --- | --- | --- | --- | --- | --- | --- | --- | --- | --- |
| **Col-0 mock control 55h** | 6,88 ± 0,31 | 12,33 ± 1,75 | 11,91 ± 0,23 | 144,94 ± 8,87 | 411,39 ± 28,53 | 810,17 ± 8,35 | 197,06 ± 5,49 | 1418,69 ± 27,01 | 454,23 ± 19,54 | 38,11 ± 0,66 | 45,58 ± 0,9 | 49,97 ± 1,74 | 16,87 ± 0,99 |
| **Col-0 auxin 55h** | 39,29 ± 1,35 | 219,19 ± 37,28 | 41,38 ± 1,85 | 382,44 ± 23,64 | 619,74 ± 16,46 | 1756,28 ± 52,95 | 239,79 ± 8,19 | 715 ± 2,65 | 531,95 ± 12,58 | 70,73 ± 2,42 | 33,83 ± 2 | 45,29 ± 0,88 | 32,09 ± 1,85 |
| **clf/cwn mock control 55h** | 9,42 ± 0,04 | 47,13 ± 14,84 | 28,91 ± 4,9 | 206,47 ± 7,93 | 530,06 ± 1,77 | 796,88 ± 53,36 | 199,15 ± 6,31 | 494,52 ± 12,84 | 607,51 ± 25,18 | 58,25 ± 1,87 | 61,93 ± 4,22 | 165,21 ± 4,4 | 40,26 ± 0,43 |
| **clf/cwn auxin 55h** | 8,06 ± 0,6 | 232,51 ± 29,18 | 28,45 ± 1,86 | 347,26 ± 20,4 | 427,59 ± 6,17 | 719,25 ± 13,7 | 166,7 ± 3,26 | 474,22 ± 43,73 | 501,94 ± 11,73 | 38,04 ± 2,56 | 32,33 ± 1,24 | 122,38 ± 6,91 | 33,71 ± 2,43 |

| **Treatment** | ***CAD*** | ***NR*** | ***ADH2*** | **Actin** | **α-Tubulin** | ***SNRK*** | ***mTOR*** | ***E2F1*** | ***E2F3*** | ***E2F5*** | **Total BAG** | **Total MC** |
| --- | --- | --- | --- | --- | --- | --- | --- | --- | --- | --- | --- | --- |
| **Col-0 mock control 55h** | 62,26 ± 0,23 | 424,57 ± 12,65 | 132,61 ± 7,72 | 794,07 ± 4,23 | 1574,36 ± 10,98 | 47,85 ± 1,11 | 11,34 ± 0,26 | 5,67 ± 0,38 | 6,45 ± 0,69 | 9,68 ± 0,45 | 228,22 ± 4,13 | 96,49 ± 2,86 |
| **Col-0 auxin 55h** | 77,96 ± 7,37 | 257,12 ± 24,49 | 163,25 ± 13,86 | 794,12 ± 26,92 | 1370,43 ± 41,2 | 46,33 ± 1,18 | 6,91 ± 0,33 | 3,82 ± 0,29 | 6,79 ± 0,54 | 14,75 ± 0,14 | 262,19 ± 6,1 | 115,22 ± 5 |
| **clf/cwn mock control 55h** | 59,08 ± 3,55 | 395,35 ± 118,08 | 159,34 ± 6,92 | 629,79 ± 14,43 | 1669,15 ± 54,96 | 57,64 ± 2,9 | 9,45 ± 0,18 | 8,23 ± 0,28 | 4,94 ± 0,28 | 10,43 ± 0,55 | 176,2 ± 3,96 | 74,45 ± 2,18 |
| **clf/cwn auxin 55h** | 39,87 ± 3,1 | 246,91 ± 39,21 | 114,75 ± 11,84 | 560,31 ± 6,91 | 1434,79 ± 49,64 | 46,93 ± 2,19 | 8,19 ± 0,58 | 7,89 ± 0,25 | 4,46 ± 0,33 | 10,57 ± 1,2 | 204,29 ± 2,79 | 58,8 ± 0,94 |

**Supplementary Table S2:** Mean values ± SE corresponding to the figure 1A. Number of biological replicates: 3. *CAD* (*cinnamyl alcohol dehydrogenase*), *NR* (*nitrate reductase*), *ADH2* (*alcohol dehydrogenase 2*), *SNRK* (*sucrose non-fermenting related kinase*), *mTOR* (*mammalian target of rapamycin*), *E2F1* (Transcription factor E2F), *BAG* (*Bcl-2 associated gene*), *MC* (*Metacaspase*).

**Supplementary Table S3:** Mean values ± SE of transcript abundance of Alternative oxidase (*AOX*) and Alcohol dehydrogenase (*ADH*) genes in *Arabidopsis* seedlings treated with auxin. The graphical representation of the values is given in figure 1A. Number of biological replicates: 3.

| **Treatment** | ***AOX1a*** | ***AOX1b*** | ***AOX1c*** | ***AOX1d*** | ***AOX2*** | **Total *AOX* (1 + 2)** | ***ADH1*** | ***ADH2*** | ***ADH* (1+2)** |
| --- | --- | --- | --- | --- | --- | --- | --- | --- | --- |
| **Col-0 mock control 55h** | 5,71 ± 0,3 | 0,02 ± 0,02 | 0,29 ± 0,08 | 0,82 ± 0,03 | 0,04 ± 0 | 6,88 ± 0,31 | 12,33 ± 1,75 | 132,61 ± 7,72 | 144,94 ± 8,87 |
| **Col-0 auxin 55h** | 17,57 ± 0,78 | 6,96 ± 0,53 | 3,12 ± 0,25 | 11,24 ± 0,42 | 0,4 ± 0,18 | 39,29 ± 1,35 | 219,19 ± 37,28 | 163,25 ± 13,86 | 382,44 ± 23,64 |
| **clf/cwn mock control 55h** | 3,25 ± 0,32 | 0 ± 0 | 0 ± 0 | 0,08 ± 0,02 | 6,09 ± 0,35 | 9,42 ± 0,04 | 47,13 ± 14,84 | 159,34 ± 6,92 | 206,47 ± 7,93 |
| **clf/cwn auxin 55h** | 2,7 ± 0,58 | 0 ± 0 | 0,02 ± 0,01 | 0 ± 0 | 5,34 ± 0,54 | 8,06 ± 0,6 | 232,51 ± 29,18 | 114,75 ± 11,84 | 347,26 ± 20,4 |

**Supplementary Table S4:** Mean values ± SE of transcript abundance of *BAG* (*Bcl-2 associated gene*) in *Arabidopsis* seedlings treated with auxin. The graphical representation of the values is given in figure 1A. Number of biological replicates: 3.

| **Treatment** | ***BAG-1*** | ***BAG-2*** | ***BAG-3*** | ***BAG-4*** | ***BAG-5*** | ***BAG-6*** | ***BAG-7*** | **Total *BAG*** |
| --- | --- | --- | --- | --- | --- | --- | --- | --- |
| **Col-0 mock control 55h** | 19,9 ± 1,76 | 3,19 ± 0,46 | 28,13 ± 1,16 | 24,72 ± 0,69 | 4,57 ± 0,18 | 2,65 ± 0,02 | 145,07 ± 1,12 | 228,22 ± 4,13 |
| **Col-0 auxin 55h** | 16,07 ± 1,64 | 1,01 ± 0,1 | 47,83 ± 4,87 | 27,19 ± 1,54 | 5,25 ± 0,27 | 1,56 ± 0,07 | 163,27 ± 0,86 | 262,19 ± 6,1 |
| **clf/cwn mock control 55h** | 8,8 ± 0,31 | 1,99 ± 0,21 | 18,6 ± 1,13 | 18,69 ± 0,82 | 5,34 ± 0,32 | 3,23 ± 0,23 | 119,54 ± 2,88 | 176,2 ± 3,96 |
| **clf/cwn auxin 55h** | 4,55 ± 0,14 | 1,18 ± 0,06 | 13,1 ± 0,71 | 19,15 ± 0,41 | 4,68 ± 0,81 | 2,24 ± 0,27 | 159,38 ± 1,96 | 204,29 ± 2,79 |

**Supplementary Table S5:** Mean values ± SE of transcript abundance of *MC* (*Metacaspase*) gene in *Arabidopsis* seedlings treated with auxin. The graphical representation of the values is given in figure 1A. Number of biological replicates: 3.

| **Treatment** | ***MC-1*** | ***MC-2*** | ***MC-3*** | ***MC-4*** | ***MC-5*** | ***MC-6*** | ***MC-7*** | ***MC-8*** | ***MC-9*** | **Total *MC*** |
| --- | --- | --- | --- | --- | --- | --- | --- | --- | --- | --- |
| **Col-0 mock control 55h** | 17,62 ± 0,57 | 0,41 ± 0,02 | 12,38 ± 0,35 | 62,24 ± 2,17 | 0,88 ± 0,18 | 0,24 ± 0,01 | 0,75 ± 0,07 | 0,07 ± 0,01 | 1,88 ± 0,22 | 96,49 ± 2,86 |
| **Col-0 auxin 55h** | 18,83 ± 0,56 | 1,96 ± 0,22 | 11,94 ± 1,19 | 73,81 ± 2,62 | 1,39 ± 0,21 | 0,52 ± 0,11 | 0,2 ± 0,03 | 0,25 ± 0,03 | 6,32 ± 0,79 | 115,22 ± 5 |
| **clf/cwn mock control 55h** | 16,67 ± 1,53 | 1,98 ± 0,13 | 4,31 ± 0,45 | 49,11 ± 0,24 | 1,44 ± 0,21 | 0,23 ± 0,09 | 0,11 ± 0,02 | 0 ± 0 | 0,61 ± 0,1 | 74,45 ± 2,18 |
| **clf/cwn auxin 55h** | 14,92 ± 0,29 | 1,65 ± 0,21 | 5,36 ± 0,84 | 35,74 ± 0,52 | 0,66 ± 0,24 | 0,13 ± 0,03 | 0 ± 0 | 0 ± 0 | 0,33 ± 0,09 | 58,8 ± 0,94 |

**Supplementary Table S6:** Mean values ± SE of transcript abundance of genes in human lung adenocarcinoma cells infected with SARS-CoV-2. The graphical representation of the values is given in figure 1E: Number of biological replicates: 3. *LDH* (*lactate dehydrogenase*), SOD1 (*superoxide dismutase1*), SOD2 (*superoxide dismutase 2*), *GPX* (*glutathione peroxidase*), *GSR* (*glutathione reductase*), *ADH5* (*alcohol dehydrogenase 5*), *PFK* (*Phosphofructokinase*), *GAPDH* (*Glyceraldehyde-3-phosphate dehydrogenase*), *G6PDH* (*glucose-6-phosphate dehydrogenase*), Cyt-MDH1 (*Cytosolic malate dehydrogenase 1*), *mt-MDH2* (*mitochondrial malate dehydrogenase*), *SNRK* (*sucrose non-fermenting related kinase*), *mTOR* (*mammalian target of rapamycin*), E2F (Transcription factor E2F), *IRF9* (*interferon regulatory factor 9*), *IRF3* (*interferon regulatory factor 3*), *NFKB1* (*nuclear factor kappa B1*), *NFKB-RelA* (*nuclear factor Kappa B- RelA*), *Caspase In* (Initiator caspases), Caspase Ex (*Executioner caspases*), *Bcl-xL*, (*B-cell lymphoma-extra large*) *ACE2* (*Angiotensin-converting enzyme 2*), *TMPRSS2* (*transmembrane protease serine 2*).

| **Treatment** | ***LDH*** | ***SOD1*** | ***SOD2*** | **Catalase** | ***GPX*** | ***GSR*** | ***ADH5*** | **Hexokinase** | ***PFK*** | ***GAPDH*** | **Enolase** | **Pyruvate Kinase** | ***G6PDH*** | ***Cyt-MDH1*** | ***mt-MDH2*** | **Actin** | **Tubulin** | ***SNRK*** | ***mTOR*** | ***E2F1*** | ***IRF9*** | ***IRF3*** | ***NF-KB1*** | ***NF-KB-RELA*** | ***Caspase In*** | ***Caspase Ex*** | ***BCL-xL*** | ***ACE2*** | ***TMPRSS2*** |
| --- | --- | --- | --- | --- | --- | --- | --- | --- | --- | --- | --- | --- | --- | --- | --- | --- | --- | --- | --- | --- | --- | --- | --- | --- | --- | --- | --- | --- | --- |
| **Mock treatment** | 1725,64 ± 72,51 | 125,13 ± 4,01 | 179,44 ± 5,07 | 33,8 ± 0,97 | 588,61 ± 27,58 | 64,95 ± 1,66 | 3,34 ± 0,68 | 17,91 ± 0,71 | 289,67 ± 2,86 | 10501,45 ± 170,44 | 1512,01 ± 73,37 | 2245,91 ± 96,04 | 750 ± 12,4 | 259,53 ± 1,14 | 170,47 ± 11,63 | 3809,94 ± 24,53 | 3102,75 ± 17,95 | 0,88 ± 0,03 | 8,69 ± 1,06 | 22,6 ± 0,89 | 8,74 ± 0,3 | 33,84 ± 2,81 | 15,63 ± 1,53 | 23,51 ± 6,52 | 19,22 ± 1,01 | 33,42 ± 1,75 | 83,41 ± 3,82 |  |  |
| **SARS-CoV-2 infected (MOI 0.2)** | 1733,98 ± 26,07 | 120,75 ± 7,77 | 196,59 ± 4,41 | 35,29 ± 1,65 | 532,36 ± 7,46 | 59,23 ± 5,34 | 3,44 ± 0,33 | 17,49 ± 0,97 | 275,94 ± 5,14 | 9532,32 ± 54,85 | 1422,75 ± 16,57 | 1970,1 ± 125,03 | 690,18 ± 9,76 | 246,94 ± 9 | 158,3 ± 5,26 | 3950,38 ± 27,56 | 2731,34 ± 91,46 | 0,71 ± 0,09 | 8,46 ± 0,77 | 20,33 ± 1,3 | 34,06 ± 1,02 | 34,58 ± 2,2 | 14,5 ± 0,55 | 26,29 ± 1,91 | 17,16 ± 0,99 | 33,51 ± 1,96 | 85,49 ± 5,21 |  |  |

**Supplementary Table S7:** Mean values ± SE of transcript abundance of genes in primary human bronchial epithelial cells infected with SARS-CoV-2. The graphical representation of the values is given in figure 1F. Number of biological replicates: 2-3. *LDH* (*lactate dehydrogenase*), SOD1 (*superoxide dismutase1*), SOD2 (*superoxide dismutase 2*), *GPX* (*glutathione peroxidase*), *GSR* (*glutathione reductase*), *ADH5* (*alcohol dehydrogenase 5*), *PFK* (*Phosphofructokinase*), *GAPDH* (*Glyceraldehyde-3-phosphate dehydrogenase*), *G6PDH* (*glucose-6-phosphate dehydrogenase*), Cyt-MDH1 (*Cytosolic malate dehydrogenase 1*), *mt-MDH2* (*mitochondrial malate dehydrogenase*), *SNRK* (*sucrose non-fermenting related kinase*), *mTOR* (*mammalian target of rapamycin*), E2F (Transcription factor E2F), *IRF9* (*interferon regulatory factor 9*), *IRF3* (*interferon regulatory factor 3*), *NFKB1* (*nuclear factor kappa B1*), *NFKB-RelA* (*nuclear factor Kappa B- RelA*), *Caspase In* (Initiator caspases), Caspase Ex (*Executioner caspases*), *Bcl-xL*, (*B-cell lymphoma-extra large*) *ACE2* (*Angiotensin-converting enzyme 2*), *TMPRSS2* (*transmembrane protease serine 2*).

| **Treatment** | ***LDH*** | ***SOD1*** | ***SOD2*** | **Catalase** | ***GPX*** | ***GSR*** | ***ADH5*** | **Hexokinase** | ***PFK*** | ***GAPDH*** | **Enolase** | **Pyruvate Kinase** | ***G6PDH*** | ***Cyt-MDH1*** | ***mt-MDH2*** | **Actin** | **Tubulin** | ***SNRK*** | ***mTOR*** | ***E2F1*** | ***IRF9*** | ***IRF3*** | ***NF-KB1*** | ***NF-KB-RELA*** | ***Caspase In*** | ***Caspase Ex*** | ***BCL-xL*** | ***ACE2*** | ***TMPRSS2*** |
| --- | --- | --- | --- | --- | --- | --- | --- | --- | --- | --- | --- | --- | --- | --- | --- | --- | --- | --- | --- | --- | --- | --- | --- | --- | --- | --- | --- | --- | --- |
| **Mock treated NHBE cells** | 1026,63 ± 15,47 | 70,14 ± 1,59 | 13,93 ± 2,65 | 17,07 ± 0,66 | 753,3 ± 31,21 | 16,57 ± 3,27 | 1,53 ± 0,23 | 50,68 ± 1,81 | 128,26 ± 6,18 | 6749,72 ± 73,27 | 447,16 ± 12,62 | 1123,24 ± 78,65 | 20,63 ± 0,25 | 70 ± 3,96 | 51,23 ± 5,07 | 348,98 ± 5,7 | 1437,39 ± 23,31 | 1,32 ± 0,07 | 5,36 ± 0,17 | 0,65 ± 0,02 | 8,67 ± 0,44 | 10,66 ± 0,7 | 16,17 ± 1 | 25,83 ± 1,06 | 7,11 ± 0,67 | 38,81 ± 0,71 | 26,33 ± 0,61 | 0,83 ± 0,09 | 2,84 ± 0,01 |
| **SARS-CoV-2 infected NHBE cells** | 1155,41 ± 3,88 | 74,33 ± 3,49 | 36,32 ± 0,33 | 14,85 ± 0,73 | 712,56 ± 25,96 | 12,72 ± 1,28 | 0,52 ± 0,06 | 51,46 ± 0,67 | 132,63 ± 1,73 | 8138,7 ± 15,11 | 496,53 ± 21,19 | 1256,24 ± 45,18 | 20,99 ± 0,15 | 61,88 ± 1,47 | 69,62 ± 0,73 | 438,19 ± 39,89 | 2161,83 ± 93,94 | 1,21 ± 0,01 | 6,04 ± 0,46 | 1,01 ± 0,02 | 20,86 ± 0,84 | 10,27 ± 1,06 | 24,4 ± 1,75 | 25,7 ± 1,54 | 8,11 ± 0,66 | 41,74 ± 1,34 | 32,84 ± 1,08 | 0,85 ± 0,1 | 2,62 ± 0,1 |
| **Mock treated A549 cells** | 1896,97 ± 87,5 | 175,42 ± 17,47 | 54,74 ± 6,91 | 38,31 ± 0,08 | 1408,57 ± 84,2 | 116,34 ± 2,06 | 4,06 ± 1,43 | 27,05 ± 1,32 | 141,82 ± 0,85 | 6835,56 ± 422,37 | 720,15 ± 4,85 | 1046,21 ± 53,43 | 548,37 ± 33,32 | 216,98 ± 6,55 | 175,43 ± 3,63 | 2928,16 ± 49,46 | 700,01 ± 2,88 | 0,97 ± 0,06 | 1,16 ± 0,2 | 11 ± 1,28 | 34,85 ± 3,43 | 47,81 ± 1,05 | 23,71 ± 0,43 | 51,29 ± 3,85 | 34,01 ± 1,75 | 32,38 ± 0,72 | 23,62 ± 2,19 | 0 ± 0 | 0 ± 0 |
| **RSV infected A549 cells** | 2282,49 ± 58,72 | 144,17 ± 35,93 | 128,73 ± 27,03 | 28,58 ± 7,44 | 1107,06 ± 35,8 | 105,85 ± 9,02 | 1,15 ± 0,31 | 35,54 ± 0,84 | 196,66 ± 21,02 | 5364,97 ± 1849,66 | 970,86 ± 0,79 | 1317,19 ± 39,83 | 495,42 ± 59,95 | 172,05 ± 42,11 | 207,31 ± 50,66 | 2870,31 ± 403,16 | 498,82 ± 73,1 | 0,52 ± 0,04 | 1,18 ± 0,21 | 6,89 ± 1,46 | 87,25 ± 1,83 | 51,32 ± 9,03 | 28,79 ± 0,41 | 64,54 ± 1,66 | 34,22 ± 5,01 | 27,26 ± 1,95 | 29,07 ± 9,87 | 0 ± 0 | 0 ± 0 |

**Supplementary Table S8:** Mean values ± SE of transcript abundance of genes in primary human lung epithelial cancer cells infected with MERS-CoV. The graphical representation of the values is given in figure 1G. Number of biological replicates: 3. *LDH* (*lactate dehydrogenase*), SOD1 (*superoxide dismutase1*), SOD2 (*superoxide dismutase 2*), *GPX* (*glutathione peroxidase*), *GSR* (*glutathione reductase*), *NOS3* (*nitric oxide synthase*), *ADH5* (*alcohol dehydrogenase 5*), *PFK* (*Phosphofructokinase*), *GAPDH* (*Glyceraldehyde-3-phosphate dehydrogenase*), *G6PDH* (*glucose-6-phosphate dehydrogenase*), Cyt-MDH1 (*Cytosolic malate dehydrogenase 1*), *mt-MDH2* (*mitochondrial malate dehydrogenase*), *SNRK* (*sucrose non-fermenting related kinase*), *mTOR* (*mammalian target of rapamycin*), E2F (Transcription factor E2F), *IRF9* (*interferon regulatory factor 9*), *IRF3* (*interferon regulatory factor 3*), *NFKB1* (*nuclear factor kappa B1*), *NFKB-RelA* (*nuclear factor Kappa B- RelA*), *Caspase In* (Initiator caspases), Caspase Ex (*Executioner caspases*), *Bcl-xL*, (*B-cell lymphoma-extra large*).

| **Treatment** | ***LDH*** | ***SOD1*** | ***SOD2*** | **Catalase** | ***GPX*** | ***GSR*** | ***NOS3*** | ***ADH5*** | **Hexokinase** | ***PFK*** | ***GAPDH*** | **Enolase** | **Pyruvate Kinase** | ***G6PDH*** | ***Cyt-MDH1*** | ***mt-MDH2*** | **Actin** | **Tubulin** | ***SNRK*** | ***mTOR*** | ***E2F1*** | ***IRF9*** | ***IRF3*** | ***NF-KB1*** | ***NF-KB-RELA*** | ***Caspase In*** | ***Caspase Ex*** | ***BCL-xL*** |
| --- | --- | --- | --- | --- | --- | --- | --- | --- | --- | --- | --- | --- | --- | --- | --- | --- | --- | --- | --- | --- | --- | --- | --- | --- | --- | --- | --- | --- |
| **Mock_3_mRNA** | 783,6 ± 16,41 | 15,44 ± 0,39 | 7,1 ± 0,32 | 9,55 ± 0,89 | 228,85 ± 2,43 | 21,15 ± 1,04 | 0,12 ± 0,02 | 1,92 ± 0,03 | 17,8 ± 1,05 | 190,32 ± 2,53 | 2560,59 ± 60 | 701,41 ± 13,97 | 1305,64 ± 23,86 | 159,97 ± 2,79 | 40,14 ± 3,31 | 111,74 ± 2,26 | 8269,19 ± 113,62 | 1413,07 ± 30,37 | 2,14 ± 0,21 | 8,25 ± 0,17 | 25,29 ± 0,59 | 1,7 ± 0,17 | 65,64 ± 1,09 | 2,03 ± 0,12 | 18,47 ± 1,07 | 10,55 ± 0,64 | 4,83 ± 0,67 | 101,56 ± 2,03 |
| **6hpi_3_mRNA** | 756,49 ± 35,21 | 12,9 ± 0,9 | 10,49 ± 0,33 | 11,4 ± 0,72 | 123,99 ± 0,68 | 15,65 ± 1,01 | 0,26 ± 0,05 | 1,61 ± 0,09 | 10,78 ± 0,99 | 107,54 ± 1,51 | 1070,49 ± 49,9 | 374,36 ± 7,27 | 723,85 ± 13,05 | 67,02 ± 1,28 | 35,62 ± 2,82 | 58,09 ± 1,33 | 4282,12 ± 124,6 | 721,07 ± 17,22 | 4,72 ± 0,19 | 9,72 ± 0,06 | 31,24 ± 1,23 | 2,71 ± 0,37 | 35,28 ± 1,54 | 2,44 ± 0,1 | 11 ± 0,34 | 8,15 ± 0,17 | 7,33 ± 0,21 | 78,34 ± 3,98 |
| **24hpi_3_mRNA** | 78 ± 3,52 | 0,4 ± 0,09 | 2,63 ± 0,37 | 1,55 ± 0,09 | 7,42 ± 1,4 | 0,78 ± 0,09 | 5,31 ± 0,09 | 0,07 ± 0 | 3,2 ± 0,17 | 11,93 ± 1,11 | 53,69 ± 4,95 | 59,39 ± 2,57 | 200,28 ± 7,17 | 3,62 ± 0,24 | 3,83 ± 0,58 | 5,13 ± 0,03 | 106,61 ± 5,12 | 33,56 ± 1,84 | 3,06 ± 0,22 | 5,77 ± 0,24 | 4,56 ± 0,33 | 4,09 ± 0,2 | 5,64 ± 0,33 | 0,93 ± 0,12 | 3,5 ± 0,54 | 1,62 ± 0,05 | 0,85 ± 0,13 | 5,66 ± 0,6 |

**Supplementary Table S9:**  Mean values ± SE of transcript abundance of genes in primary human fetal lung fibroblast cells infected with MERS-CoV. The graphical representation of the values is given in figure 1H. Number of biological replicates: 3. *LDH* (*lactate dehydrogenase*), SOD1 (*superoxide dismutase1*), SOD2 (*superoxide dismutase 2*), *GPX* (*glutathione peroxidase*), *GSR* (*glutathione reductase*), *ADH5* (*alcohol dehydrogenase 5*), *PFK* (*Phosphofructokinase*), *GAPDH* (*Glyceraldehyde-3-phosphate dehydrogenase*), *G6PDH* (*glucose-6-phosphate dehydrogenase*), Cyt-MDH1 (*Cytosolic malate dehydrogenase 1*), *mt-MDH2* (*mitochondrial malate dehydrogenase*), *SNRK* (*sucrose non-fermenting related kinase*), *mTOR* (*mammalian target of rapamycin*), E2F (Transcription factor E2F), *IRF9* (*interferon regulatory factor 9*), *IRF3* (*interferon regulatory factor 3*), *NFKB1* (*nuclear factor kappa B1*), *NFKB-RelA* (*nuclear factor Kappa B- RelA*), *Caspase In* (Initiator caspases), Caspase Ex (*Executioner caspases*), *Bcl-xL*, (*B-cell lymphoma-extra large*).

| **Treatment** | ***LDH*** | ***SOD1*** | ***SOD2*** | **Catalase** | ***GPX*** | ***GSR*** | ***ADH5*** | **Hexokinase** | ***PFK*** | ***GAPDH*** | **Enolase** | **Pyruvate Kinase** | ***G6PDH*** | ***Cyt-MDH1*** | ***mt-MDH2*** | **Actin** | **Tubulin** | ***SNRK*** | ***mTOR*** | ***E2F1*** | ***IRF9*** | ***IRF3*** | ***NF-KB1*** | ***NF-KB-RELA*** | ***Caspase In*** | ***Caspase Ex*** | ***BCL-xL*** |
| --- | --- | --- | --- | --- | --- | --- | --- | --- | --- | --- | --- | --- | --- | --- | --- | --- | --- | --- | --- | --- | --- | --- | --- | --- | --- | --- | --- |
| **MOCK-MRC5-3** | 445,32 ± 21,79 | 32,44 ± 1,38 | 21,23 ± 1,95 | 10,89 ± 0,59 | 171,47 ± 13,57 | 9,81 ± 0,57 | 11,14 ± 0,15 | 21,66 ± 0,39 | 106,25 ± 1,46 | 1443,08 ± 42,01 | 504,98 ± 29,44 | 322,49 ± 6,55 | 46,77 ± 1,11 | 43,36 ± 0,94 | 69,88 ± 1,33 | 6310,98 ± 123,72 | 634,71 ± 47,37 | 3,48 ± 0,09 | 5,08 ± 0,17 | 9,26 ± 0,21 | 4,63 ± 0,31 | 10,4 ± 0,36 | 6,05 ± 0,61 | 14,63 ± 0,35 | 3,41 ± 0,12 | 17,17 ± 2,09 | 9,76 ± 0,45 |
| **VMERS_MERS-MRC5lowMOI-24hr-3** | 388,97 ± 5,7 | 26,52 ± 0,47 | 15,36 ± 0,65 | 8,94 ± 0,36 | 127,86 ± 5,6 | 7,63 ± 0,62 | 9,09 ± 0,53 | 18,23 ± 0,78 | 93,15 ± 4,9 | 1167,03 ± 38,58 | 457,55 ± 14,22 | 281,91 ± 9,16 | 34,27 ± 2,11 | 35,55 ± 0,93 | 56,1 ± 2,69 | 5436,71 ± 312,34 | 604,06 ± 14,57 | 3,18 ± 0,35 | 4,8 ± 0,36 | 8,57 ± 0,55 | 4,05 ± 0,1 | 9,68 ± 0,39 | 5,66 ± 0,07 | 14,01 ± 0,54 | 2,85 ± 0,12 | 13,43 ± 1,57 | 8,87 ± 0,45 |
| **VMERS_MERS-MRC5lowMOI-48hr-3** | 95,07 ± 2,67 | 4,36 ± 0,35 | 6,12 ± 0,51 | 2,04 ± 0,05 | 19,15 ± 1,2 | 1,66 ± 0,06 | 1,15 ± 0,15 | 5,24 ± 0,15 | 19,77 ± 2,37 | 150,98 ± 13,85 | 100,43 ± 5,08 | 81,94 ± 9,61 | 3,63 ± 0,29 | 5,29 ± 0,22 | 10,98 ± 0,82 | 652,84 ± 10,17 | 77,28 ± 4,7 | 3,01 ± 0,12 | 3,28 ± 0,19 | 2,34 ± 0,04 | 1,95 ± 0,23 | 2,55 ± 0,14 | 5,18 ± 0,25 | 4,84 ± 0,26 | 0,93 ± 0,03 | 5,34 ± 0,25 | 3,84 ± 0,3 |
| **VMERS_MERS-MRC5HighMOI-24hr-3** | 230,77 ± 1,1 | 12,17 ± 0,23 | 9,61 ± 0,82 | 4,52 ± 0,31 | 55,32 ± 3,92 | 4,28 ± 0,3 | 4,29 ± 0,07 | 9,88 ± 0,24 | 50,66 ± 1,14 | 530,96 ± 6,93 | 259,59 ± 10,86 | 159,22 ± 2,01 | 14,11 ± 0,76 | 15,45 ± 0,5 | 29,75 ± 0,8 | 2762,83 ± 59,06 | 299,75 ± 6,75 | 2,87 ± 0,1 | 3,52 ± 0,12 | 5,77 ± 0,05 | 2,67 ± 0,3 | 6,02 ± 0,23 | 4,44 ± 0,18 | 8,28 ± 0,24 | 2,16 ± 0,13 | 7,22 ± 0,54 | 6,03 ± 0,14 |
| **VMERS_MERS-MRC5HighMOI-48hr-3** | 131,08 ± 3,46 | 5,8 ± 0,19 | 8,61 ± 0,57 | 2,23 ± 0,16 | 24,28 ± 0,88 | 2,38 ± 0,33 | 1,69 ± 0,13 | 7,14 ± 0,51 | 24,59 ± 0,17 | 213,17 ± 9,74 | 129,28 ± 2,61 | 83,61 ± 0,7 | 4,71 ± 0,29 | 7,54 ± 0,38 | 13,45 ± 0,49 | 721,76 ± 14,53 | 116,93 ± 0,92 | 2,95 ± 0,16 | 4,06 ± 0,16 | 3,63 ± 0,11 | 2,09 ± 0,17 | 2,97 ± 0,23 | 7,33 ± 0,71 | 5,23 ± 0,73 | 1,26 ± 0,05 | 5,61 ± 0,31 | 4,58 ± 0,35 |

| **Treatment** | ***LDH*** | ***SOD1*** | ***SOD2*** | **Catalase** | ***GPX*** | ***GSR*** | ***ADH5*** | **Hexokinase** | ***PFK*** | ***GAPDH*** | **Enolase** | **Pyruvate Kinase** | ***G6PDH*** | ***Cyt-MDH1*** | ***mt-MDH2*** | **Actin** | **Tubulin** | ***SNRK*** | ***mTOR*** | ***E2F1*** | ***IRF9*** | ***IRF3*** | ***NF-KB1*** | ***NF-KB-RELA*** | ***Caspase In*** | ***Caspase Ex*** | ***BCL-xL*** |
| --- | --- | --- | --- | --- | --- | --- | --- | --- | --- | --- | --- | --- | --- | --- | --- | --- | --- | --- | --- | --- | --- | --- | --- | --- | --- | --- | --- |
| **MOCK-MRC5-3** | 445,32 ± 21,79 | 32,44 ± 1,38 | 21,23 ± 1,95 | 10,89 ± 0,59 | 171,47 ± 13,57 | 9,81 ± 0,57 | 11,14 ± 0,15 | 21,66 ± 0,39 | 106,25 ± 1,46 | 1443,08 ± 42,01 | 504,98 ± 29,44 | 322,49 ± 6,55 | 46,77 ± 1,11 | 43,36 ± 0,94 | 69,88 ± 1,33 | 6310,98 ± 123,72 | 634,71 ± 47,37 | 3,48 ± 0,09 | 5,08 ± 0,17 | 9,26 ± 0,21 | 4,63 ± 0,31 | 10,4 ± 0,36 | 6,05 ± 0,61 | 14,63 ± 0,35 | 3,41 ± 0,12 | 17,17 ± 2,09 | 9,76 ± 0,45 |
| **VMERS_SARS-MRC5lowMOI-24hr-3** | 428,37 ± 5,41 | 25,85 ± 1,08 | 17,43 ± 0,67 | 10,66 ± 0,31 | 138,85 ± 6,16 | 9,02 ± 0,29 | 10,36 ± 0,12 | 19,54 ± 1,01 | 96,73 ± 3,18 | 1152,66 ± 2,42 | 478,75 ± 24,31 | 296,85 ± 13,84 | 41,28 ± 0,77 | 36,84 ± 1,14 | 58,79 ± 0,68 | 5733,41 ± 132,02 | 559,28 ± 30,08 | 3,94 ± 0,18 | 5,87 ± 0,31 | 9,18 ± 0,68 | 4,04 ± 0,46 | 11,44 ± 0,84 | 5,13 ± 0,3 | 14,56 ± 0,33 | 3,62 ± 0,18 | 17,76 ± 1,45 | 9,58 ± 0,31 |
| **VMERS_SARS-MRC5lowMOI-48hr-3** | 485,51 ± 19,91 | 29,22 ± 2,6 | 30,23 ± 0,8 | 9,7 ± 1,41 | 156,4 ± 13,71 | 8,94 ± 1,38 | 12,25 ± 1,69 | 16,1 ± 0 | 81,65 ± 6,66 | 1230,13 ± 82,05 | 407,01 ± 39,72 | 251,8 ± 5,76 | 37,36 ± 5,88 | 38,99 ± 0,7 | 58,1 ± 5,75 | 4254,71 ± 228,15 | 433,07 ± 12,38 | 4,06 ± 0,39 | 4,89 ± 0,15 | 4,72 ± 0,68 | 4,18 ± 0,54 | 8,15 ± 0,47 | 5,34 ± 0,7 | 12,08 ± 0,06 | 3,58 ± 0,56 | 17,79 ± 0,23 | 6,81 ± 0,45 |
| **VMERS_SARS-MRC5HighMOI-24hr-3** | 391,37 ± 14,08 | 25,81 ± 0,28 | 15,92 ± 1,28 | 10,77 ± 0,63 | 125,15 ± 5,14 | 8,85 ± 0,14 | 9,3 ± 0,18 | 17,57 ± 1,56 | 86,85 ± 3,48 | 1147,26 ± 34,08 | 293,34 ± 5,13 | 321,38 ± 4,94 | 39,39 ± 1,75 | 42,85 ± 1,42 | 61,04 ± 2,6 | 6217,12 ± 213,81 | 586,48 ± 20,7 | 3,93 ± 0,23 | 5,77 ± 0,24 | 8,94 ± 0,13 | 4,96 ± 0,39 | 11,07 ± 0,25 | 4,88 ± 0,22 | 15,36 ± 0,13 | 3,55 ± 0,63 | 18,24 ± 0,53 | 9,4 ± 0,7 |
| **VMERS_SARS-MRC5HighMOI-48hr-3** | 455,23 ± 25,58 | 30,45 ± 3,04 | 22,99 ± 2,47 | 9,67 ± 0,45 | 178,91 ± 9,51 | 9,21 ± 0,79 | 12,19 ± 0,68 | 20,94 ± 1,57 | 84,87 ± 4,22 | 1218,48 ± 120,38 | 409,97 ± 1,78 | 307,09 ± 38,61 | 44,15 ± 0,76 | 38,88 ± 2,37 | 64,17 ± 5,87 | 5253,34 ± 178,02 | 460,24 ± 9,93 | 4,39 ± 0,11 | 4,7 ± 0,3 | 4,99 ± 0,34 | 4,33 ± 0,47 | 9,62 ± 0,99 | 5,57 ± 0,02 | 16,16 ± 1,36 | 3,75 ± 0,71 | 23,72 ± 2,29 | 8,92 ± 0,53 |

**Supplementary Table S10:**  Mean values ± SE of transcript abundance of genes in primary human fetal lung fibroblast cells infected with SARS-CoV. The graphical representation of the values is given in figure 1H. Number of biological replicates: 2-3. *LDH* (*lactate dehydrogenase*), SOD1 (*superoxide dismutase1*), SOD2 (*superoxide dismutase 2*), *GPX* (*glutathione peroxidase*), *GSR* (*glutathione reductase*), *ADH5* (*alcohol dehydrogenase 5*), *PFK* (*Phosphofructokinase*), *GAPDH* (*Glyceraldehyde-3-phosphate dehydrogenase*), *G6PDH* (*glucose-6-phosphate dehydrogenase*), Cyt-MDH1 (*Cytosolic malate dehydrogenase 1*), *mt-MDH2* (*mitochondrial malate dehydrogenase*), *SNRK* (*sucrose non-fermenting related kinase*), *mTOR* (*mammalian target of rapamycin*), E2F (Transcription factor E2F), *IRF9* (*interferon regulatory factor 9*), *IRF3* (*interferon regulatory factor 3*), *NFKB1* (*nuclear factor kappa B1*), *NFKB-RelA* (*nuclear factor Kappa B- RelA*), *Caspase In* (Initiator caspases), Caspase Ex (*Executioner caspases*), *Bcl-xL*, (*B-cell lymphoma-extra large*).

**Supplementary Table S11:** Bioproject details of RNA-seq experiments used to obtain the gene expression in *Arabidopsis thaliana* and Human cells

| BioProject  acession | Species | Physiological stage/ treatment | Experiment details | Tissue/Cell type | Biological  replicates | Reference |
| --- | --- | --- | --- | --- | --- | --- |
| PRJNA320769 | *Arabidopsis thaliana* | Somatic embryogenesis | The RNA seq analyses were performed in *Arabidopsis* wt (Col-0 genotype) and clf/swn mutant. Expression profiles of wt and clf/swn were analyzed in control and auxin treatment at time 55h.  Seeds from WT and mutant clf/swn genotypes were placed as complete seedlings without wounding for 55h into *in vitro*-culture medium, which contained 5μM 2,4-D and 1% sucrose (Mozgová et al., 2017). Under these conditions, WT seedlings induced non-embryogenic callus, while the mutant induced embryo formation. Mozgová et al. (2017) highlighted that overall performance of the mutant indicated accelerated SE-induction due to depletion of polycomb repressive complex 2 (PRC2): if immature, WT zygotic seeds were SE-competent when treated with 2,4-D; however, 3d of treatment were required to obtain only around 10% of SE-forming zygotic embryos. To reach about 70% success rate, 7d were required. In the mutant, 60hrs of 2,4-D-treatment were already sufficient to induce SE in 87% of shoot explants, whereas WT shoots were non-SE-competent (Mozgová et al. 2017). Furthermore, the authors highlighted that biological processes activated in the mutant in response to the SE-inductive treatment were mainly related to oxidative stress response and cell wall remodeling. | stem | Three | Mozgová  et al. 2017 |
| PRJNA615032 | *Homo sapiens* | SARS-CoV-2 infection (MOI 2) | Primary human lung epithelium (NHBE) were mock treated or infected with SARS-CoV-2 (USA-WA1/2020). Time point: 24hrs after treatment. | Cell line: NHBE;  Cell type: primary human bronchial epithelial cells | Three | Blanco-Melo et al., 2020 |
|  |  | SARS-CoV-2 infection (MOI 0.2) | Transformed lung alveolar (A549) cells were mock treated or infected with SARS-CoV-2 (USA-WA1/2020). Time point: 24hrs after treatment. | Cell line: A549;  Cell type: Lung adenocarcinoma | Three |  |
|  |  | RSV infection (MOI 15) | A549 cells were mock treated or infected with respiratory syncytial vírus (RSV) A2 strain. Time point: 24hrs after treatment. | Cell line: A549;  Cell type: Lung adenocarcinoma | Two |  |
| PRJNA580021 | *Homo sapiens* | MERS-CoV infection (MOI 4) | human lung adenocarcinoma epithelial (Calu-3) cells were mock-infected or infected with MERS-CoV. mRNA analyses were performed at times 6 and 24 h after treatment. | Cell line: Calu-3;  Cell type: Lung adenocarcinoma | Three | Zhang et al., 2020 |
| PRJNA233943 | *Homo sapiens* | MERS-CoV infection (0.1 and 3 MOI) | MRC5 cells were infected at MOI of 0.1 and 3 with MERS-CoV and RNA was isolated at 24 and 48 hours post infection. | Cell line: MRC-5 cells  Cell type: [fibroblasts](https://en.wikipedia.org/wiki/Fibroblast) of lung tissue | At least 3 Biological replicates and technical replicates | Data are published only in SRA database from GenBank (NCBI) |
|  |  | SARS-CoV infection (0.1 and 3 MOI) | MRC5 cells were infected at MOI of 0.1 and 3 with SARS-CoV and RNA was isolated at 24 and 48 hours post infection. | Cell line: MRC-5 cells  Cell type: [fibroblasts](https://en.wikipedia.org/wiki/Fibroblast) of lung tissue |  |  |
