## Supplementary Figures S1 and S2 for "ROS/RNS balancing, aerobic fermentation regulation and cell cycle control – a complex early trait (‘CoV-MAC-TED’) for combating SARS-CoV-2-induced cell reprogramming"

**Supplementary Figure S1**: Differences in transcript levels (normalized in RPKM values) of AOX, ADH1, LDH and Enolase in WT and clf/cwn-mutant between mock controls and between 55h auxin-treated seeds. **: highly significant (p < 0.01) differences

**

**

**

**

**

AOX

|  | Total AOX |
| --- | --- |
| Col-0 mock | 6.88 |
| clf/cwn mock | 9.42 |
| Col-0 auxin | 39.29 |
| clf/cwn auxin | 8.06 |

|  | ADH1 |
| --- | --- |
| Col-0 mock | 12.33 |
| clf/cwn mock | 47.13 |
| Col-0 auxin | 219.19 |
| clf/cwn auxin | 232.51 |

|  | LDH |
| --- | --- |
| Col-0 mock | 11.91 |
| clf/cwn mock | 28.91 |
| Col-0 auxin | 41.38 |
| clf/cwn auxin | 28.45 |

|  | Enolase |
| --- | --- |
| Col-0 mock | 411.39 |
| clf/cwn mock | 530.06 |
| Col-0 auxin | 619.74 |
| clf/cwn auxin | 427.59 |

**Supplementary Figure S2**: Differences in transcript levels (normalized in RPKM) of AOX1 and AOX2 between mock control and auxin-treated seeds of *Arabidopsis thaliana* (**: highly significant (p < 0.01))

|  | AOX2 |
| --- | --- |
| Col-0 mock | 0.176839 |
| Col-0 auxin | 0.403902 |
| clf/cwn mock | 6.08701 |
| clf/cwn auxin | 5.34338 |

|  | AOX1 |
| --- | --- |
| Col-0 mock | 6.87706 |
| Col-0 auxin | 38.9324 |
| clf/cwn mock | 3.34194 |
| clf/cwn auxin | 2.73001 |
